# Ankyrin-B regulates cardiomyocyte sarcomere organization and its protein-protein interactions are significantly altered by p.S646F expression

**DOI:** 10.64898/2026.10.29.762332

**Authors:** Nicole S. York, K’sana Wood Lynes-Ford, Alexa C.H. McAdam, Melissa Paradis, Hope Maron, Connor A. Larkey, Leigh E. Wicki-Stordeur, Laura T. Arbour, Leigh Anne Swayne

**Affiliations:** School of Medical Sciences, Faculty of Health, University of Victoria, Victoria, British Columbia, Canada; Department of Medical Genetics, University of British Columbia, Victoria, British Columbia, Canada; Island Medical Program, University of British Columbia, Victoria, British Columbia, Canada

**Author notes:** Correspondence should be addressed to Leigh Anne Swayne.

**Keywords:** Ankyrin-B, *ANK2*, cardiomyocytes, protein interaction networks, α-actinin-1, cardiomyopathy, sarcomere

## Abstract

Ankyrin-B (ANKB) is a large (220 kDa) multi-domain membrane-localized cytoplasmic protein that acts as a scaffold coordinating cytoskeletal, ion channel, and transporter proteins critical for proper cardiomyocyte structure and function. There is a prevalence of structural cardiac abnormalities within the cohort of the novel ANKB p.S646F variant that we previously identified in two multigenerational Gitxsan First Nation families, suggesting a role for ANKB in cardiac structure. To investigate the impact of ANKB on cardiomyocyte structural organization, we imaged sarcomere structure in primary cardiomyocytes from cardiac-specific ANKB heterozygous (expressing lower levels of ANKB) and full knock-out mice using confocal microscopy. We observed marked disruption of sarcomere structure in primary cardiac cultures associated with lower ANKB expression, suggesting ANKB is required for proper development of sarcomeres. To shed light on the underlying mechanisms, we next employed an unbiased proteomics approach to identify novel ANKB protein-protein interactions. This analysis uncovered a novel ANKB interaction with α-actinin-1, an actin-binding protein that stabilizes focal adhesions essential for development of cardiomyocyte structure. Western blot comparison of α-actinin-1 co-precipitation with wildtype ANKB and ANKB p.S646F, revealed increased α-actinin-1 enrichment with ANKB p.S646F. Bioinformatics analysis of the ANKB interactome revealed enrichment of cytoskeletal organization, sarcomere structure, and chaperone proteins, and marked changes with ANKB p.S646F. Together findings point to a novel role for ANKB in the regulation of cardiomyocyte structural development and support additional investigation into a link between p.S646F and cardiac structural abnormalities.

## Introduction

We have previously identified a variant in the *ANK2* gene, encoding for the ankyrin-B (ANKB) scaffold protein, ANKB p.S646F - the first reported loss-of-function variant within ANKB’s membrane binding domain responsible for interacting with ion flux proteins - associated with a range of cardiac functional and structural abnormalities [1]. This discovery expanded a growing list of ANKB variants associated with functional (e.g. arrhythmias) and structural (cardiomyopathy) cardiac phenotypes, with the former (e.g. long QT syndrome (LQTS) type 4) [2–5] having received relatively more focus of mechanistic study (reviewed in [6,7]). The physical interaction of ANKB with ion flux proteins bolstered investigation of ANKB variants relative to function, albeit not without controversy [8]. These flux proteins include sodium potassium ATPase (NKA), sodium calcium exchanger (NCX), inositol 1,4,5-trisphosphate receptor (IP3R), and the Kir6.2 inwardly-rectifying ATP-sensitive potassium channel pore subunit [9,10]. ANKB also physically interacts with phosphatase 2A (PP2A), which regulates the phosphorylation state of ryanodine receptor 2 (RYR2) [11], the channel primarily responsible for the increase in cytoplasmic Ca^2+^ associated with contraction. Conversely, the link between ANKB variants and cardiac structural phenotypes has been relatively understudied. However, ANKB physically interacts with cytoskeletal components involved in membrane organization, sarcomere integrity, and cardiomyocyte differentiation (reviewed in [7]), like βII-spectrin, obscurin, dystrophin, dystroglycan, and β-catenin [5,12–14]; therefore ANKB is well-positioned to regulate cardiomyocyte shape, and it is therefore perhaps not so surprising that ANKB p.S646F is associated with cardiac structural phenotypes. This work aims to investigate the molecular and cellular basis of the relationship between ANKB p.S646F and cardiac structural phenotypes.

Our initial characterization of the ANKB p.S646F variant included 18 individuals across two multigeneration Gitxsan First Nation families [1]. These individuals were initially identified in the context of known LQTS within the community but were negative for the potassium voltage-gated channel subfamily Q member 1 (KCNQ1) variants that typically account for the high prevalence of LQTS within this population. Importantly, the phenotype in those with the ANKB p.S646F variant included arrhythmias, seizures, cardiac structural malformations, and sudden cardiac death [1]. Our updated analysis of the expanded cohort of individuals with the p.S646F variant, presented here, supports the prevalence of cardiac structural phenotypes, underscoring a potential link between the p.S646F variant and cardiomyocyte structure.

Little is known about the role of ANKB in regulating cardiomyocyte structure, but there is good reason to suggest it could. Our prior investigations of the cellular impact of the p.S646F variant revealed reduced expression levels compared to wild type (WT) ANKB, suggesting instability and/or increased degradation in the H9c2 cardiomyoblast cell line, and mis- localization (along with NCX) in primary cardiomyocytes [1,15]. H9c2 cells expressing ANKB p.S646F exhibited increased cell death and proliferation [15]. Together with our observation that retinoic-acid differentiation of H9c2 cells triggered an increase in ANKB expression levels [16], and ANKB physical interaction with cytoskeletal proteins, it is reasonable to suspect that ANKB plays a role in the structural development of cardiomyocytes. If ANKB p.S646F affected that role, this could shed light on the pathophysiology of the associated cardiac structural abnormalities. However, key gaps in understanding, such as the impact of the loss of ANKB on cardiomyocyte structure have not been addressed. Moreover, our insight into the ANKB interactome in cardiomyocytes is limited to targeted studies and the impact of the ANKB p.S646F variant on the ANKB interactome is unknown.

To that end, we first determined the impact of ANKB on sarcomere organization as a critical readout of cardiomyocyte structure. Reduced ANKB expression (conditional cardiac partial and full *Ank2* KO mice) in primary cardiomyocytes disrupted sarcomere organization, supporting ANKB’s role in structural organization. We then investigated the ANKB cardiac interactome, using unbiased methods. With immunoprecipitation assays from the H9c2 cell line (to achieve appropriate levels of protein abundance for proper proteomic identification), coupled to liquid chromatography and tandem mass spectrometry (LC-MS), and downstream bioinformatics analyses, we found that the ANKB interactome is enriched in cytoskeletal organization and sarcomere structure proteins. We identified a novel interaction between ANKB and α-actinin-1, a critical protein that promotes adhesion enlargement required for sarcomere assembly during cardiomyocyte structural development, that is enriched by ANKB p.S646F.

In summary, we present updated clinical findings on an expanded cohort of ANKB p.S646F carriers, confirming the presence of structural cardiac abnormalities in this population. We show that partial loss of ANKB expression markedly disrupts sarcomere organization in primary cardiomyocytes. Finally, we report the first cardiac ANKB interactome, revealing interactions with cytoskeleton elements, and its potential shift with ANKB p.S646F. Together our clinical and cellular data suggest a prominent role for ANKB in regulating cardiomyocyte structure and support its consideration in future clinical and mechanistic studies.

## Results

### Updated clinical findings in 68 individuals with the ANKB p.S646F variant show increased structural cardiac abnormalities

We previously reported a novel *ANK2* c.1937C>T variant (ANKB p.S646F) inherited in an autosomal dominant manner, in two multigenerational Gitxsan families, identified in the context of LQTS, and associated with arrhythmia, seizure, syncope, and structural heart disease [1]. At that time, we found 18 individuals heterozygous for the p.S646F variant. Our cohort now includes a total of 68 ANKB p.S646F heterozygous carriers (59 adults and 9 children) among 740 participants, including individuals identified through cascade screening and broader participant testing. Of these, 66 were tested for the two common LQTS-associated variants in Northern British Columbia First Nations populations, KCNQ1 p.V205M and p.R591H [17], with 7 of the 66 tested participants also carrying the p.V205M variant. Here, we present an update on the clinical findings since 2017, summarized in Table 1.

**Table 1.**
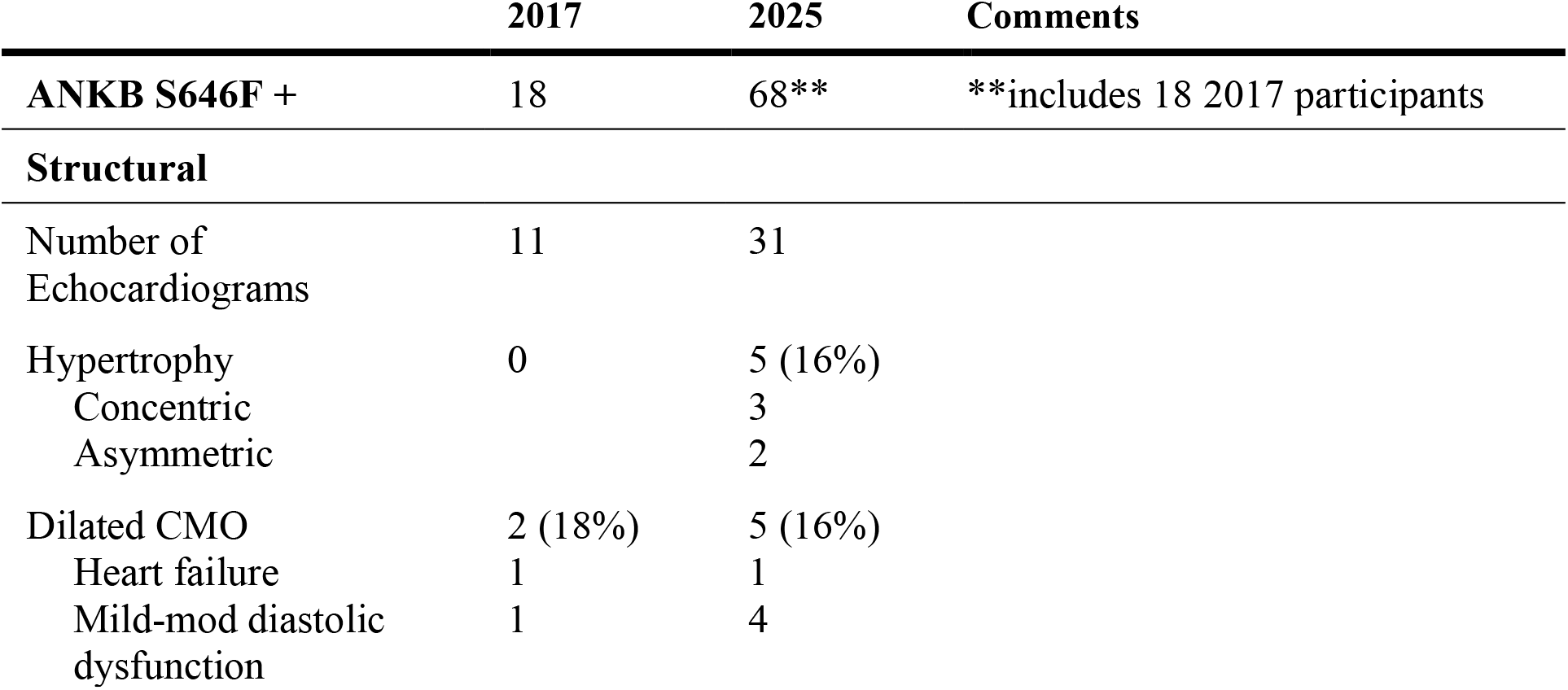
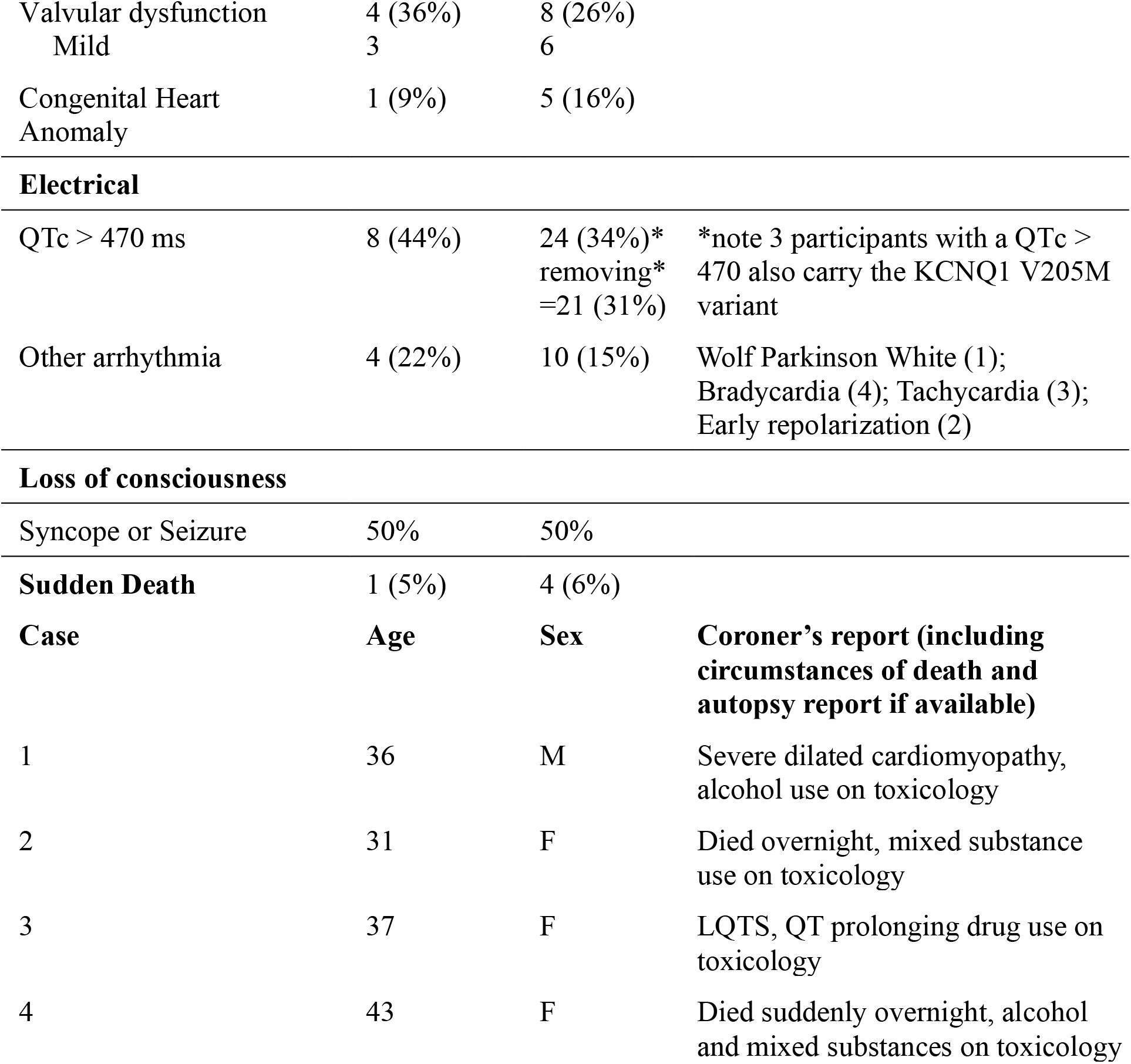
ANKB S646F clinical features: 2025 update since 2017.

|  | 2017 | 2025 | Comments |
| --- | --- | --- | --- |
| **ANKB S646F +** | 18 | 68** | **includes 18 2017 participants |
| **Structural** |  |  |  |
| Number of Echocardiograms | 11 | 31 |  |
| Hypertrophy | 0 | 5 (16%) |  |
| Concentric |  | 3 |  |
| Asymmetric |  | 2 |  |
| Dilated CMO | 2 (18%) | 5 (16%) |  |
| Heart failure | 1 | 1 |  |
| Mild-mod diastolic dysfunction | 1 | 4 |  |
| Valvular dysfunction | 4 (36%) | 8 (26%) |  |
| Mild | 3 | 6 |  |
| Congenital Heart Anomaly | 1 (9%) | 5 (16%) |  |
| **Electrical** |  |  |  |
| QTc > 470 ms | 8 (44%) | 24 (34%)*<br>removing*<br>=21 (31%) | *note 3 participants with a QTc > 470 also carry the KCNQ1 V205M variant |
| Other arrhythmia | 4 (22%) | 10 (15%) | Wolf Parkinson White (1);<br>Bradycardia (4); Tachycardia (3);<br>Early repolarization (2) |
| **Loss of consciousness** |  |  |  |
| Syncope or Seizure | 50% | 50% |  |
| **Sudden Death** | 1 (5%) | 4 (6%) |  |
| **Case** | **Age** | **Sex** | **Coroner's report (including circumstances of death and autopsy report if available)** |
| 1 | 36 | M | Severe dilated cardiomyopathy, alcohol use on toxicology |
| 2 | 31 | F | Died overnight, mixed substance use on toxicology |
| 3 | 37 | F | LQTS, QT prolonging drug use on toxicology |
| 4 | 43 | F | Died suddenly overnight, alcohol and mixed substances on toxicology |

Among the 68 identified with the ANKB p.S646F variant, 64 have had at least one ECG, 31 have undergone echocardiography, and 34 have been evaluated through the adult or pediatric BC Inherited Arrhythmia Program (BCIAP) clinics, where multidisciplinary genetic and heart rhythm specialist assessment occurs [18]. Evaluation of the expanded cohort continues to demonstrate structural cardiac abnormalities, (cardiomyopathy/congenital heart anomalies) in those with echocardiogram reports. In addition, a greater number of sudden deaths relative to the original report were observed. Structural cardiac abnormalities observed included concentric (3/31) and asymmetric (2/31) hypertrophy (16%, 5/31); dilated cardiomyopathy (16%, 5/31), heart failure (1/31), and congenital heart anomalies (16%, 5/31). Of note, and consistent with our previous report, there was minimal evidence of QTc prolongation in individuals under 25 years but, for those over age 25 years (31% exhibited a prolonged QTc (excluding those with the p.V205M variant). Beyond the prolonged QT interval, heart rhythm variation was documented (15%), including Wolff-Parkinson-White syndrome, bradycardia, tachycardia, and early repolarization patterns. Those with the variant experienced frequent loss of consciousness episodes such as syncope or seizure, occurring in half of carriers, consistent with the initial findings. There have been four sudden deaths (6%), all in young adults, documented among p.S646F carriers, three since the first report. Autopsy reports revealed severe dilated cardiomyopathy in one case (previously reported), overnight deaths with mixed substance use on toxicology (2 cases), and LQTS with QT-prolonging drug use on toxicology (1 case). These findings support the association of ANKB p.S646F with diverse cardiac manifestations, including structural abnormalities, heart rhythm disturbance, and sudden death.

### ANKB loss disrupts sarcomere organization in primary cardiomyocyte cultures

The prevalence of structural cardiac abnormalities in individuals with ANKB p.S646F, coupled with our previous cellular results demonstrating reduced expression of the p.S646F variant in H9c2 cells [1,15], led us to investigate whether ANKB expression levels impact cardiomyocyte structure. To this end, we generated primary cardiomyocyte cultures from male and female control, and cardiac-specific ANKB heterozygous (HET) and ANKB knock-out (KO) mice. Western blotting of primary cardiomyocyte lysates confirmed decreased ANKB expression in HET cultures and almost complete loss in KO cultures (Figure 1A). A two-way ANOVA was performed with sex and genotype as the main factors (Figure 1B). There was a significant main effect of genotype (F(3,16) = 48.70, p < 0.0001), but no significant main effect of sex (F(1,16) = 0.03, p = 0.8652), and no significant sex x genotype interaction (F(3,16) = 0.09, p = 0.9641). Šídák’s multiple comparisons test revealed significant ANKB reduction in female HET cultures compared to control (female control: 100.0% ± 6.5%, female HET: 51.4% ± 3.5%, p = 0.0108), and drastic loss in female KO cultures compared to controls (female KO: 5.9% ± 1.9%, p < 0.0001). Similarly, ANKB levels were significantly reduced in male HET cultures compared to controls (male control: 100.0% ± 13.8%, male HET: 58.2% ± 8.2%, p = 0.0322), and male KO cultures showed almost complete loss of ANKB relative to controls (male KO: 3.6% ± 0.2%, p < 0.0001).

**Figure 1.**
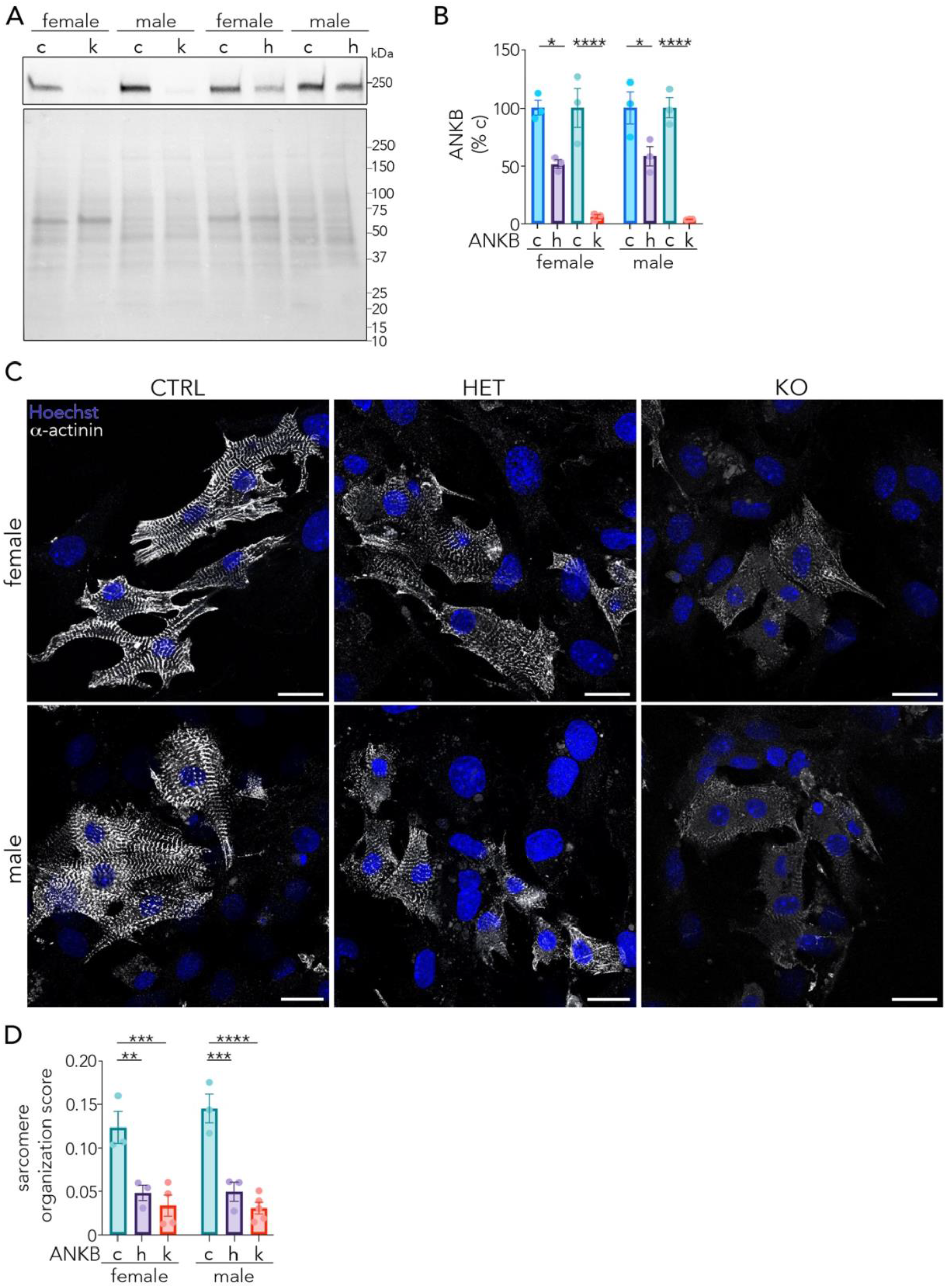
ANKB loss disrupts sarcomere organization in primary cardiomyocyte cultures. (A) Representative images of wildtype (WT, w), heterozygous (HET, h), and ANKB knockout (KO, k) cultures stained for α-actinin (white) and Hoechst (blue). WT cultures displayed organized sarcomeric α-actinin structure, while HET and KO cultures showed significant loss of sarcomere organization in both female and male cultures. (B) Quantification of sarcomere organization score was significantly reduced in both HET and KO cultures compared with WT controls in both female and male conditions. (C) Representative Western blot of ANKB protein expression in WT, HET, and KO cultures from female and male samples. (D) Densitometry analyses of ANKB protein expression demonstrated reduced ANKB abundance in HET cultures and near-complete loss in KO cultures in both sexes. Immunocytochemistry: N = 3 independent cultures for WT and HET female and male conditions, N = 4 for female KO cultures, and N = 5 for male KO cultures. Western blot: N = 3 independent biological replicates. Lysates were run independently on separate gels between replicates and quantified per replicate. ANKB was normalized to total protein and expressed as percent control (WT). Data are presented as mean ± SEM. WT = blue or teal, HET = purple, KO = red. ****, p < 0.0001; ***, p < 0.001; **, p < 0.01; *, p < 0.05; unlabeled, p > 0.05 by two-way ANOVA and Šidák’s multiple comparisons test. See Tables 2 and 3 for detailed statistical analysis. Scale bar = 20 µm.

**Table 2.** Summary statistics of protein expression analysis in primary cardiomyocyte cultures. Descriptive statistics including minimum (Min) and maximum (Max) values for each condition, as well as the mean ± SEM, are listed. Two-way ANOVA was used to assess the effects of genotype (CTRL, HET, and KO), sex (female and male), and the genotype × sex interaction on ANKB expression levels. N = 3 independent biological replicates. Šidák’s multiple comparisons test was used for post hoc comparisons between genotypes within each sex and between sexes within each genotype. Mean differences and 95% confidence intervals (CI) are reported for each comparison. ****, *p* < 0.0001; ***, *p* < 0.001; **, *p* < 0.01; *, *p* < 0.05; ns, not significant (*p* > 0.05).

| descriptive statistics |  |  |  |  |
| --- | --- | --- | --- | --- |
| sex | ANKB | min (%) | max (%) | normalized percent to WT |
| female | CTRL | 90.8 | 112.5 | 100 ± 6.5 |
|  | HET | 44.9 | 56.7 | 51.4 ± 3.5 |
|  | CTRL | 68.8 | 126.7 | 100 ± 16.9 |
|  | KO | 2.2 | 8.5 | 5.9 ± 1.9 |
| male | CTRL | 74.6 | 121.9 | 100 ± 13.8 |
|  | HET | 44.1 | 72.7 | 58.2 ± 8.2 |
|  | CTRL | 85.9 | 116.2 | 100 ± 8.8 |
|  | KO | 3.2 | 3.8 | 3.6 ± 0.2 |
| 2- way ANOVA |  |  |  |  |
| Effect | F (df1, df2) | p-value |  |  |
| Genotype | F (3, 16) = 48.66 | p < 0.0001 |  |  |
| Sex | F (1, 16) = 0.0298 | p = 0.8652 |  |  |
| Genotype x Sex | F (3, 16) = 0.0907 | p = 0.9641 |  |  |
| Šidák's multiple comparisons test |  |  |  |  |
| Comparison | ANKB | Mean Diff | 95% CI | p-value |
| female | CTRL | 48.57 | 9.560 to 87.59 | 0.0108 |
|  | HET |  |  | * |
|  | CTRL | 94.07 | 55.06 to 133.1 | <0.0001 |
|  | KO |  |  | **** |
| male | CTRL | 41.80 | 2.786 to 80.81 | 0.0322 |
|  | HET |  |  | * |
|  | CTRL | 96.36 | 57.34 to 135.4 | <0.0001 |
|  | KO |  |  | **** |
| HET | female | -6.774 | -43.26 to 29.71 | 0.9768 |
|  | male |  |  | ns |
| KO | female | 2.285 | -34.20 to 38.77 | 0.9996 |
|  | male |  |  | ns |

**Table 3.** Summary statistics of sarcomere organization analysis in primary cardiomyocyte cultures. Two-way ANOVA was used to assess the effects of genotype (WT, HET, and KO), sex (female and male), and the genotype × sex interaction on sarcomere organization. Šidák’s multiple comparisons test was used for post hoc comparisons between genotypes within each sex and between sexes within each genotype. Immunocytochemistry analyses included N = 3 independent cultures for CTRL and HET female and male conditions, N = 4 for female KO cultures, and N = 5 for male KO cultures. 10 cells across 4 ROIs were analyzed per N. Mean differences and 95% confidence intervals (CI) are reported for each comparison. ****, *p* < 0.0001; ***, *p* < 0.001; **, *p* < 0.01; *, *p* < 0.05; ns, not significant (*p* > 0.05).

| 2- way ANOVA |  |  |  |  |
| --- | --- | --- | --- | --- |
| Effect | F (df1, df2) | p-value |  |  |
| Genotype | F (2, 15) = 40.00 | p < 0.0001 |  |  |
| Sex | F (1, 15) = 0.4635 | p = 0.5064 |  |  |
| Genotype x Sex | F (2, 15) = 0.5772 | p = 0.5735 |  |  |
| Šidák's multiple comparisons test |  |  |  |  |
| Comparison | Mean Diff | Mean Diff | 95% CI | p-value |
| female | WT | 0.07525 | 0.02629 to 0.1242 | 0.0027 |
|  | HET |  |  | ** |
|  | WT | 0.0898 | 0.04400 to 0.1356 | 0.0003 |
|  | KO |  |  | *** |
|  | HET | 0.01456 | -0.03124 to 0.06036 | 0.7913 |
|  | KO |  |  | ns |
| male | WT | 0.0956 | 0.04663 to 0.1446 | 0.0003 |
|  | HET |  |  | *** |
|  | WT | 0.1144 | 0.07060 to 0.1582 | <0.0001 |
|  | KO |  |  | **** |
|  | HET | 0.01879 | -0.02500 to 0.06259 | 0.6065 |
|  | KO |  |  | ns |
| WT | female | -0.02175 | -0.07071 to 0.02722 | 0.5807 |
|  | male |  |  | ns |
| HET | female | -0.001397 |  | 0.9998 |
|  | male |  | -0.05036 to<br>0.04756 | ns |
| KO | female<br>male | 0.00284 | -0.03739 to<br>0.04307 | 0.9968<br>ns |

We next assessed sarcomere organization as a readout of cardiomyocyte structure in control, HET, and KO cultures. We performed immunostaining and confocal imaging of α-actinin-2 to visualize sarcomeres (Figure 1C), followed by quantification using the SarcOmere Texture Analysis (SOTA) method [19] in which a score of ≥0.1 indicates organized sarcomere structure. Control cultures exhibited SOTA scores indicative of organized sarcomere structure (0.1232 ± 0.0183 female; 0.1450 ± 0.0158 male), while HET and KO cultures had significantly lower scores (0.0480 ± 0.0089 female, 0.0494 ± 0.0111 male, and 0.0334 ± 0.0120 female, 0.0306 ± 0.0065 male, respectively). A two-way ANOVA was performed with sex and genotype as factors. There was a significant main effect of genotype (F(2,15) = 40.00, p < 0.0001), but no significant main effect of sex (F(1,15) = 0.46, p = 0.5064). There was no significant sex x genotype interaction (F(2,15) = 0.58, p = 0.5735). Šídák’s multiple comparisons test revealed significantly reduced sarcomere organization in female HET (p = 0.0027) and KO (p = 0.0003) cultures compared with female controls, and in male HET (p = 0.0003) and KO (p < 0.0001) cultures compared with male controls (Figure 1D). Together, our findings demonstrate that reductions in ANKB expression disrupt typical cardiomyocyte sarcomere organization.

### ANKB forms a novel interaction with α-actinin-1 that is enhanced by the p.S646F variant

To gain insight into potential mechanisms underlying the role of ANKB in regulating cardiomyocyte structure, we next explored the cardiac ANKB interactome. Using the rat H9c2 cardiomyoblast cell line, we performed GFP immunoprecipitations from cells transfected with GFP-tagged wildtype (WT) ANKB, ANKB p.S646F variant, or GFP alone, then analyzed these samples by liquid chromatography coupled to tandem mass spectrometry (LC-MS). All protein hits identified in the GFP alone control samples were excluded from ANKB WT and ANKB p.S646F protein lists.

Our unbiased screen identified a potential novel interaction between ANKB and α-actinin-1, a protein required for focal adhesion enlargement, which links the developing sarcomere to the extracellular matrix, and is emerging as a regulator of sarcomere assembly [20]. Given our results implicated ANKB in sarcomere organization, we next validated this novel ANKB-α-actinin-1 interaction and assessed whether it was modulated by p.S646F variant expression. Western blotting of GFP immunoprecipitations from H9c2 cells expressing GFP-tagged ANKB WT, ANKB p.S646F, or GFP alone confirmed enrichment of α-actinin-1 in ANKB lanes compared to the GFP negative control (Figure 2 A). Notably, enrichment of α-actinin-1 was approximately doubled with expression of the p.S646F variant compared to ANKB WT (WT = 100% ± 13.1%, S646F = 196.5% ± 32.1%, p = 0.0498, unpaired T-test, Figure 2 Bi).

**Figure 2.**
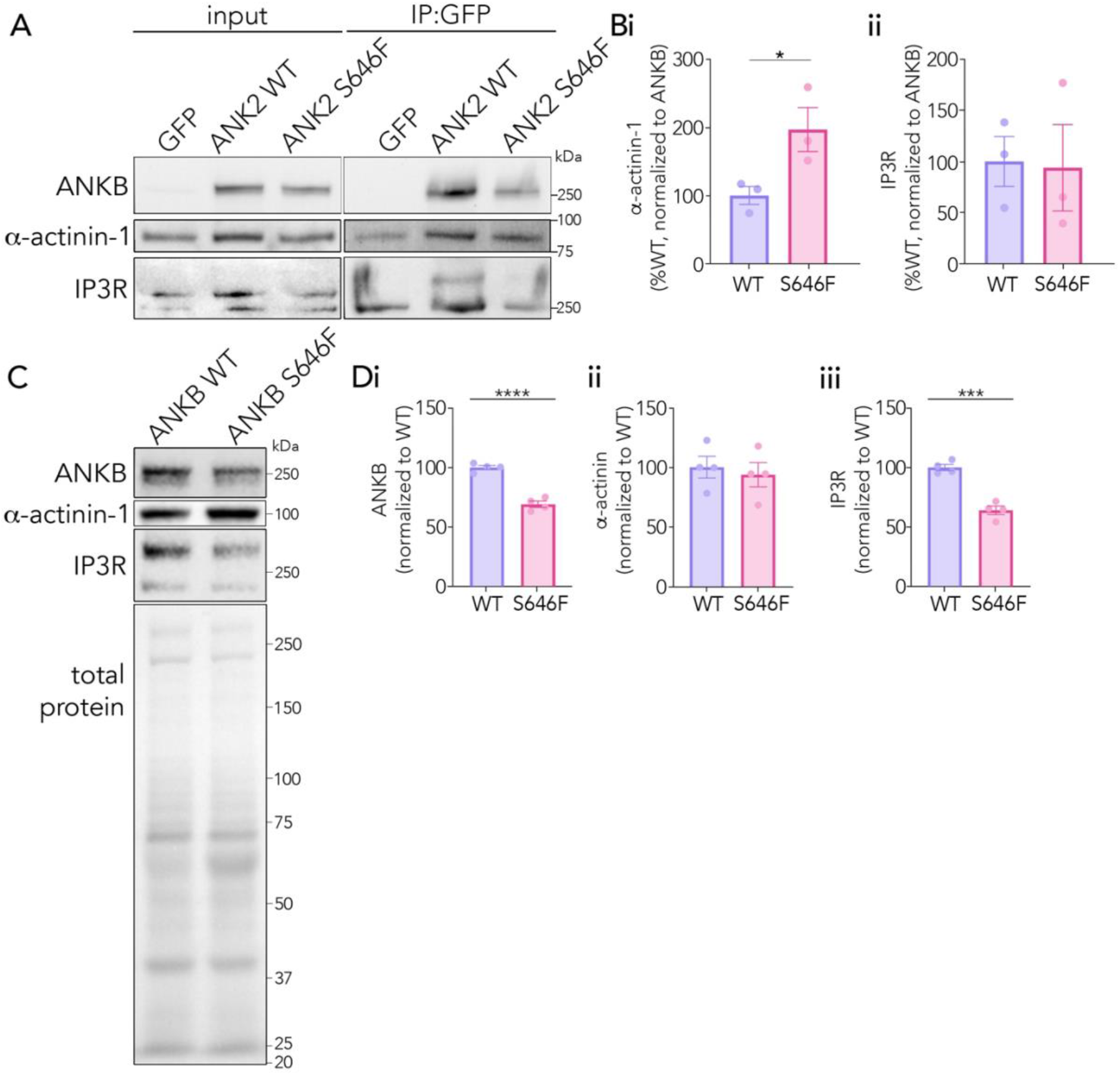
ANKB interacts with α-actinin-1 and IP3R in H9c2 cells. (A) Representative Western blots of GFP-immunoprecipitations and inputs from H9c2 cells transiently expressing ANKB WT-GFP, ANKB S646F-GFP or GFP alone (control), and probed for ANKB, α-actinin, and IP3R. (Bi) Densitometry analysis revealed an enrichment in α-actinin-1 signal with ANKB S646F-GFP compared to ANKB WT-GFP (WT = 100% ± 13.1%, S646F = 196.5% ± 32.1%, p=0.0498). (Bii) Densitometry analysis of IP3R was not significantly different between ANKB S646F-GFP and ANKB WT-GFP (WT = 100% ± 24.4%, S646F = 93.9% ± 42.2%, p = 0.9057). N = 3 independent biological replicates (passages). α-actinin and IP3R signals were normalized to ANKB-GFP levels for each condition and expressed as percent WT. (C) Representative Western blots of whole cell lysates from ANKB WT or S646F transfected H9c2 cells. (C) Densitometry analyses revealed (i) ANKB protein expression was significantly reduced in S646F-transfected cells compared with WT (WT = 100 ± 1.8%, S646F = 69.2 ± 2.7%, p = 0.0002) (ii) α-actinin-1 expression was not significantly different between groups (WT = 100 ± 9.0%, S646F = 93.7 ± 10.1%, p = 0.6576) (iii) IP3R expression was significantly reduced in S646F-transfected cells compared with WT (WT = 100 ± 2.6%, S646F = 64.0 ± 3.6%, p = 0.0003). N = 4 independent biological replicates (passages). Normalized to total protein and expressed as percent WT. Lysates were run independently on separate gels between replicates and quantified per replicate. Data is presented as mean ± SEM. WT = purple, S646F = magenta. ****, p < 0.0001; ***, p < 0.001; **, p < 0.01; *, p < 0.05; unlabeled, p > 0.05 by unpaired t test. See Table 4 for detailed statistical analysis.

**Table 4.** Summary statistics of protein expression analysis in H9c2 cells transfected with ANKB WT or p.S646F for immunoprecipitation and whole cell lysate experiments. Mean ± SEM values for each condition are listed. For immunoprecipitation experiments, N = 3 independent biological replicates (passages). α-actinin-1 and IP3R signals were normalized to ANKB-GFP levels for each condition and expressed as percent WT. For whole cell lysate experiments, N = 4 independent biological replicates (passages). ANKB, α-actinin-1, and IP3R signals were normalized to total protein and expressed as percent WT. Lysates were run independently on separate gels between replicates and quantified per replicate. Unpaired t-tests were used to compare ANKB WT and S646F conditions. ****, p < 0.0001; ***, p < 0.001; **, p < 0.01; *, p < 0.05; ns, not significant (p > 0.05).

| protein | ANKB | normalized<br>percent to WT | p-value unpaired<br>T-test | t(df) |
| --- | --- | --- | --- | --- |
| **immunoprecipitation** |  |  |  |  |
| α-actinin | WT | 100 ± 13.1 | 0.0498 | t(4) = 2.78 |
| | S646F | 196.5 ± 32.1 | * | |
| IP3R | WT | 100 ± 24.4 | 0.9057 | t(4) = 0.13 |
| | S646F | 93.4 ± 42.2 | ns | |
| **whole cell lysates** |  |  |  |  |
| ANKB | WT | 100 ± 1.8 | <0.0001 | t(6) = 9.57 |
| | S646F | 69.2 ± 2.7 | **** | |
| α-actinin | WT | 100 ± 9.0 | 0.6574 | t(6) = 0.47 |
| | S646F | 93.7 ± 10.1 | ns | |
| IP3R | WT | 100 ± 2.6 | 0.0002 | t(6) = 8.12 |
| | S646F | 64.0 ± 3.6 | *** | |

We also confirmed an ANKB-IP3R interaction within our system, which was previously identified in cardiomyocytes [21]. Intriguingly, although the IP3R interaction was earlier mapped to the ANKB membrane-binding domain, we saw no significant difference in enrichment of IP3R between ANKB WT and p.S646F conditions (WT = 100% ± 24.4%, S646F = 93.9% ± 42.2%, p = 0.9057, unpaired T-test, Figure 2 Bii). Finally, we used Western blotting of H9c2 cells to investigate whole cell expression levels of ANKB, α-actinin-1 and IP3R when ANKB WT or p.S646F variant were expressed (Figure 2 C). Consistent with our previous findings [1,15], we confirmed ANKB p.S646F expression was significantly reduced compared to ANKB WT (WT = 100 ± 1.8, S646F = 69.2 ± 2.7, unpaired T-test, p < 0.0001, Figure 2 Di).

Furthermore, while α-actinin-1 levels remained unchanged between WT and p.S646F conditions (WT = 100 ± 9.0, S646F = 93.7 ± 10.1, unpaired T-test, p = 0.6574, Figure 2 Dii), IP3R expression was significantly reduced (WT = 100 ± 2.6, S646F = 64.0 ± 3.6, unpaired T-test, p = 0.0002, Figure 2 Diii).

### The ANKB interactome consists of cytoskeletal, sarcomere, chaperone, and metabolic proteins and is altered by p.S646F

To further elucidate the molecular functions and pathways associated with ANKB and its role in cardiac structural phenotypes, we performed complementary bioinformatic analyses of our unbiased proteomics data. Together, our proteomics screen identified 23 and 68 potential interactors from ANKB WT and p.S646F conditions, respectively, with 8 of these shared between the two. Qualitatively, the ANKB p.S646F variant was associated with a greater number of potential interactors than WT, suggesting differences in the interactome profiles between the two conditions. However, since mass spectrometry is abundance-biased [22,23], it is possible that proteins identified in a single group may have been present in both but escaped detection.

For instance, we originally identified α-actinin-1 in the p.S646F group with the proteomics screen. Yet, Western blot validation showed that α-actinin-1 co-immunoprecipitated with both ANKB WT and p.S646F, while remaining significantly enriched with the p.S646F variant (Figure 2 A,B). Therefore, we combined WT and p.S646F protein lists to use in downstream bioinformatics analyses, yielding a combined ANKB interactome of 83 unique protein hits. We first performed Search Tool for the Retrieval of Interacting Genes/Proteins (STRING) network analysis [24,25], to identify existing functional relationships between our putative ANKB interacting proteins (Figure 3 A). STRING analysis revealed functional clustering of proteins in cytoskeleton (Figure 3 B), sarcomere (Figure 3 C), chaperone (Figure 3 D), and histone-associated networks.

**Figure 3.**
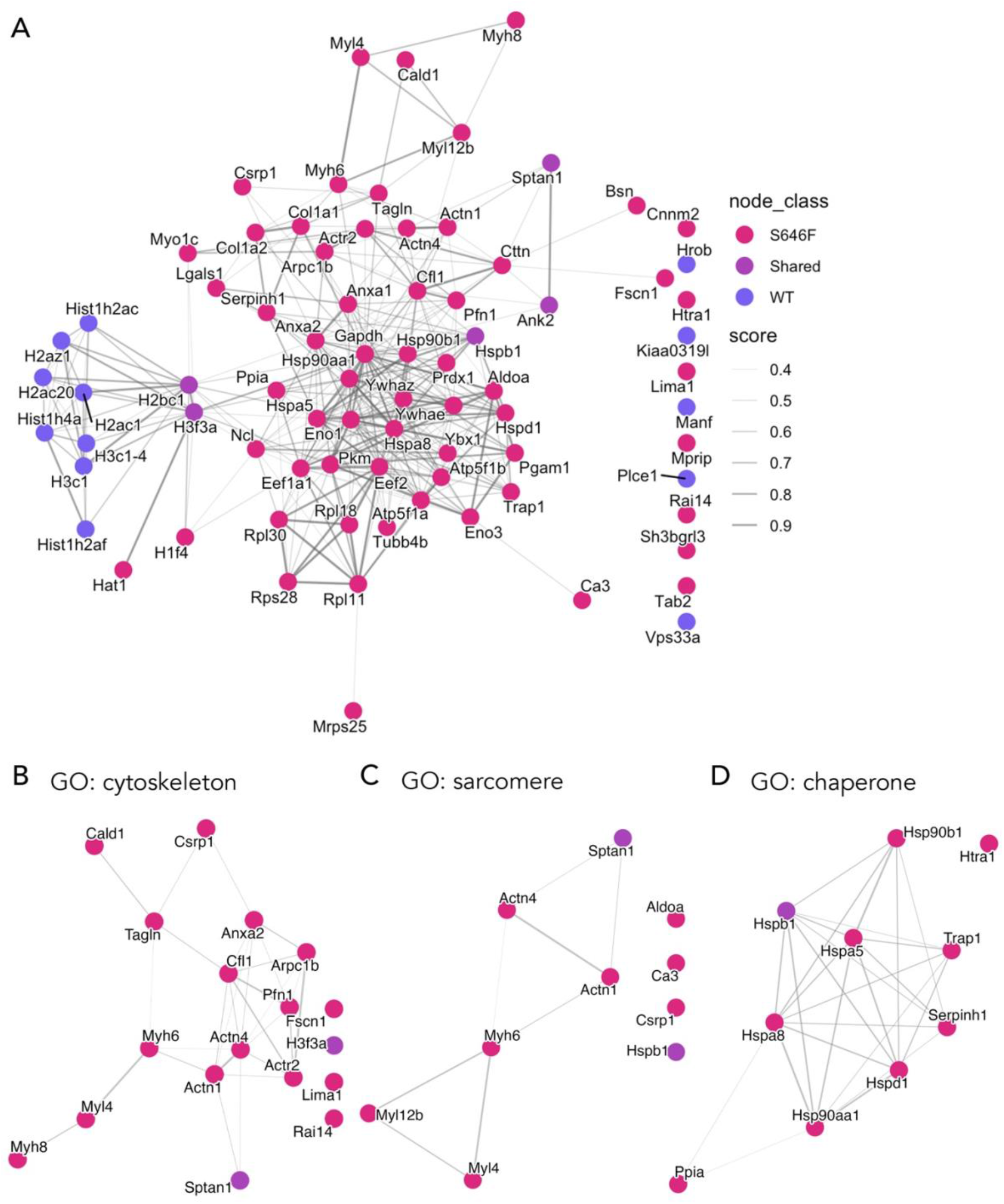
STRING network analysis identifies cytoskeleton-, sarcomeric-, and chaperone-associated proteins within the ANKB interactome. (A) STRING protein interaction network generated from the merged WT and ANKB p.S646F interactomes identified via immunoprecipitation followed by proteomics. STRING analysis identified enrichment of proteins associated with (B) cytoskeletal structure, (C) sarcomeric organization, and (D) chaperone activity. Proteins identified uniquely in WT (purple), uniquely in S646F (magenta), or shared between conditions (lavender) are indicated by node color. Edge thickness represents the confidence score of predicted protein–protein associations within the STRING network.

We next used the PANTHER classification system to identify prevalent Gene Ontology (GO) [26,27] Biological Process and Molecular Function terms, and PANTHER Protein Class terms within our ANKB interactome. In the Biological Process category, prevalent GO terms included cellular process (GO:0009987), biological regulation (GO:0065007), metabolic process (GO:0008152), localization (GO:0051179), response to stimulus (GO:0050896), and developmental process (GO:0032502) (Figure 4 A). For Molecular Function, prominent GO terms were binding (GO:0005488), catalytic activity (GO:0003824), and structural molecule activity (GO:0005198) (Figure 4 B). Common PANTHER Protein Class identifiers were cytoskeletal proteins (PC00085), chromatin/chromatin-binding or -regulatory proteins (PC00077), translational proteins (PC00263), metabolite interconversion enzymes (PC00262), and chaperones (PC00072) within the interactome (Figure 4 C). We subsequently performed PANTHER overrepresentation analyses to quantify GO and PANTHER terms overrepresented within our ANKB interactome. Full summaries are presented in Tables 4–8. Differentially regulated terms with an FDR <0.05 were considered significant. Within the GO Biological Process category, key overrepresented terms included response to heat (GO:0009408; +46.48 fold enrichment, FDR = 0.0252), heterochromatin organization (GO:0070828; +43.28 fold enrichment, FDR = 4.07 x 10^-10^), actin filament bundle organization (GO:0061572; +32.18 fold enrichment, FDR = 0.00462), and muscle cell development (GO:0055001; +20.92 fold enrichment, FDR = 0.0139), among others, and broadly fell into clusters associated with gene and chromatin regulation, cytoskeleton, metabolic processes, and muscle development (Table 5). In the GO Molecular Function category, overrepresented terms included nucleosome binding (GO:0031490; +41.63 fold enrichment, FDR = 6.33 x 10^-12^) and actin binding (GO:0003779; +18.59 fold enrichment, FDR = 8.24 x 10^-13^), and fell into broader clusters associated with chromatin binding, actin binding, unfolded protein binding, and lyase activity (Table 6). In the GO Cellular Component category, significantly overrepresented terms included actin cytoskeleton (GO:0015629; +18.59 fold enrichment, FDR = 1.44 x 10^-13^), sarcomere (GO:0030017; +12.68 fold enrichment, FDR = 0.0312), and chromatin (GO:0000785; +8.45 fold enrichment, FDR = 8.29 x 10^-5^), broadly falling into clusters related to cytoskeleton, contractile fiber, and chromatin (Table 7). Overrepresented PANTHER Protein Class terms included HSP90 family chaperone (PC00028; +>100 fold enrichment, FDR = 4.34×10^-5^), HSP70 family chaperone (PC00027; +39.84 fold enrichment, FDR = 0.0221), actin or actin-binding cytoskeletal protein (PC00041; +16.64 fold enrichment, FDR = 4.03 x 10^-14^), and chromatin/chromatin-binding or -regulatory protein (PC00077; +12.79 fold enrichment, FDR = 6.41 x 10^-11^) (Table 8). Overrepresented PANTHER Pathway terms included Glycolysis (P00024; +19.92 fold enrichment, FDR = 0.00263), Cytoskeletal regulation by Rho GTPase (P00016; +17.01 fold enrichment, FDR = 0.000927), and Integrin signalling pathway (P00034; +7.38 fold enrichment, FDR = 0.016) (Table 9).

**Figure 4.**
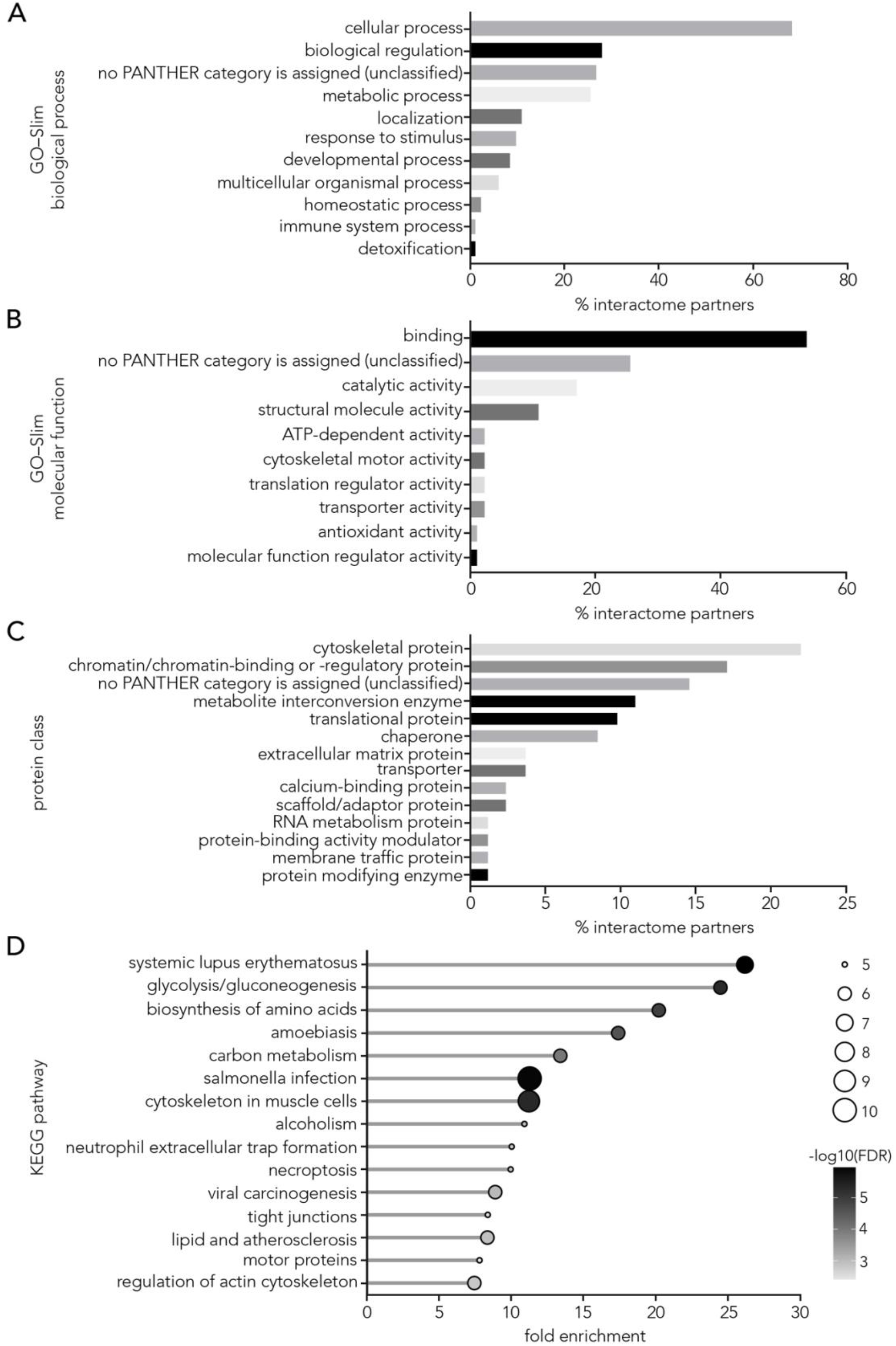
GO-Slim functional classification and KEGG pathway enrichment of the ANKB interactome. (A) GO-Slim biological process analysis identified enrichment of proteins associated with cellular process, biological regulation, metabolic process, response to stimulus, and localization. (B) GO-Slim molecular function analysis identified enrichment of proteins involved in binding, catalytic activity, and structural molecule activity. (C) Protein class analysis identified enrichment of cytoskeletal proteins, translational proteins, chromatin/chromatin-binding or -regulatory proteins, metabolite interconversion enzymes, and chaperones within the interactome. (D) KEGG pathway enrichment analysis identified pathways associated with cytoskeletal organization and cellular metabolism, including regulation of the actin cytoskeleton, cytoskeleton in muscle cells, glycolysis/gluconeogenesis, and carbon metabolism. In panels A–C, percentages represent the proportion of proteins within the merged ANKB interactome assigned to each category. In panel D, bubble size represents gene count and bubble color represents −log10(FDR).

**Table 5.** PANTHER overrepresentation analysis of Biological Process GO terms represented in the ANKB interactome. Fold enrichment is calculated from the observed number of genes/proteins in the ANKB interactome (# interactome) divided by the expected number of hits (expected), based on the total number of genes within the *Rattus norvegicus* genome classified under the same GO term (# genome). Enrichment was considered significant with a false discovery rate (FDR) < 0.05. Row shading indicates GO terms within a hierarchical set.

| PANTHER GO-Slim Biological Process | GO ID | # genome | # interactome | expected | fold enrichment | FDR |
| --- | --- | --- | --- | --- | --- | --- |
| response to temperature stimulus | GO:0009266 | 16 | 2 | 0.06 | 34.86 | 0.0429 |
| response to heat | GO:0009408 | 12 | 2 | 0.04 | 46.48 | 0.0252 |
| epigenetic regulation of gene expression | GO:0040029 | 80 | 9 | 0.29 | 31.38 | 4.16E-09 |
| negative regulation of biosynthetic process | GO:0009890 | 439 | 9 | 1.57 | 5.72 | 0.0014 |
| negative regulation of cellular process | GO:0048523 | 1013 | 12 | 3.63 | 3.3 | 0.00926 |
| negative regulation of cellular metabolic process | GO:0031324 | 547 | 10 | 1.96 | 5.1 | 0.00133 |
| negative regulation of cellular biosynthetic process | GO:0031327 | 439 | 9 | 1.57 | 5.72 | 0.00137 |
| negative regulation of biological process | GO:0048519 | 1127 | 13 | 4.04 | 3.22 | 0.0071 |
| negative regulation of metabolic process | GO:0009892 | 611 | 11 | 2.19 | 5.02 | 0.000688 |
| negative regulation of macromolecule metabolic process | GO:0060255 | 591 | 11 | 2.12 | 5.19 | 0.000551 |
| negative regulation of macromolecule biosynthetic process | GO:0010558 | 435 | 9 | 1.56 | 5.77 | 0.00137 |
| negative regulation of gene expression | GO:0010629 | 195 | 9 | 0.7 | 12.87 | 4.8E-06 |
| negative regulation of gene expression, epigenetic | GO:0045814 | 64 | 9 | 0.23 | 39.22 | 6.21E-10 |
| cellular process | GO:0009987 | 9025 | 58 | 32.36 | 1.79 | 1.94E-06 |
| cellular component organization or biogenesis | GO:0071840 | 2471 | 34 | 8.86 | 3.84 | 4.11E-10 |
| cellular component organization | GO:0016043 | 2289 | 34 | 8.21 | 4.14 | 1.81E-10 |
| protein-containing complex organization | GO:0043933 | 554 | 13 | 1.99 | 6.54 | 0.000011 |
| protein-DNA complex organization | GO:0071824 | 215 | 10 | 0.77 | 12.97 | 0.000001 |
| chromatin organization | GO:0006325 | 189 | 10 | 0.68 | 14.76 | 3.17E-07 |
| chromatin remodeling | GO:0006338 | 172 | 10 | 0.62 | 16.22 | 1.41E-07 |
| heterochromatin organization | GO:0070828 | 58 | 9 | 0.21 | 43.28 | 4.07E-10 |
| heterochromatin formation | GO:0031507 | 54 | 9 | 0.19 | 46.48 | 3.08E-10 |
| organelle organization | GO:0006996 | 1381 | 17 | 4.95 | 3.43 | 0.000479 |
| cytoskeleton organization | GO:0007010 | 604 | 14 | 2.17 | 6.46 | 4.41E-06 |
| actin filament-based process | GO:0030029 | 284 | 12 | 1.02 | 11.78 | 1.08E-07 |
| actin cytoskeleton organization | GO:0030036 | 272 | 12 | 0.98 | 12.3 | 7.54E-08 |
| supramolecular fiber organization | GO:0097435 | 307 | 11 | 1.1 | 9.99 | 2.09E-06 |
| actin filament organization | GO:0007015 | 161 | 8 | 0.58 | 13.86 | 1.45E-05 |
| Arp2/3 complex-mediated actin nucleation | GO:0034314 | 15 | 2 | 0.05 | 37.19 | 0.0381 |
| actin filament bundle organization | GO:0061572 | 26 | 3 | 0.09 | 32.18 | 0.00462 |
| actin filament bundle assembly | GO:0051017 | 24 | 3 | 0.09 | 34.86 | 0.00376 |
| pyridine-containing compound metabolic process | GO:0072520 | 91 | 5 | 0.33 | 15.32 | 0.00104 |
| pyridine nucleotide metabolic process | GO:0019362 | 89 | 5 | 0.32 | 15.67 | 0.000984 |
| nicotinamide nucleotide metabolic process | GO:0046496 | 89 | 5 | 0.32 | 15.67 | 0.00101 |
| carbohydrate metabolic process | GO:0005975 | 154 | 5 | 0.55 | 9.06 | 0.00883 |
| carbohydrate catabolic process | GO:0016052 | 81 | 5 | 0.29 | 17.22 | 0.000676 |
| ribonucleoside triphosphate metabolic process | GO:0009199 | 92 | 6 | 0.33 | 18.19 | 9.94E-05 |
| nucleoside triphosphate metabolic process | GO:0009141 | 102 | 6 | 0.37 | 16.41 | 0.000174 |
| purine nucleoside triphosphate metabolic process | GO:0009144 | 89 | 6 | 0.32 | 18.8 | 8.58E-05 |
| purine ribonucleoside triphosphate metabolic process | GO:0009205 | 88 | 6 | 0.32 | 19.02 | 8.44E-05 |
| ATP metabolic process | GO:0046034 | 86 | 6 | 0.31 | 19.46 | 7.78E-05 |
| purine ribonucleotide metabolic process | GO:0009152 | 172 | 6 | 0.62 | 9.73 | 0.00174 |
| ribose phosphate metabolic process | GO:0019693 | 193 | 6 | 0.69 | 8.67 | 0.0031 |
| ribonucleotide metabolic process | GO:0009259 | 181 | 6 | 0.65 | 9.25 | 0.00222 |
| carbohydrate derivative catabolic process | GO:1901136 | 129 | 5 | 0.46 | 10.81 | 0.00441 |
| ribonucleotide catabolic process | GO:0009264 | 70 | 5 | 0.25 | 19.92 | 0.000431 |
| nucleotide catabolic process | GO:0009166 | 80 | 5 | 0.29 | 17.43 | 0.000655 |
| nucleotide metabolic process | GO:0009117 | 233 | 6 | 0.84 | 7.18 | 0.00756 |
| purine nucleotide metabolic process | GO:0006163 | 210 | 6 | 0.75 | 7.97 | 0.00455 |
| purine-containing compound metabolic process | GO:0072521 | 224 | 6 | 0.8 | 7.47 | 0.00634 |
| purine-containing compound catabolic process | GO:0072524 | 74 | 5 | 0.27 | 18.84 | 0.000491 |
| purine nucleotide catabolic process | GO:0009160 | 71 | 5 | 0.25 | 19.64 | 0.000445 |
| purine ribonucleotide catabolic process | GO:0009150 | 68 | 5 | 0.24 | 20.51 | 0.000404 |
| generation of precursor metabolites and energy | GO:0006091 | 163 | 5 | 0.58 | 8.56 | 0.0109 |
| nucleoside diphosphate metabolic process | GO:0009132 | 72 | 5 | 0.26 | 19.37 | 0.00046 |
| nucleobase-containing small molecule metabolic process | GO:0055086 | 278 | 6 | 1 | 6.02 | 0.0169 |
| nucleoside phosphate metabolic process | GO:0006753 | 243 | 6 | 0.87 | 6.89 | 0.00898 |
| organic cyclic compound catabolic process | GO:1901361 | 207 | 5 | 0.74 | 6.74 | 0.0272 |
| cellular nitrogen compound catabolic process | GO:0044270 | 190 | 5 | 0.68 | 7.34 | 0.0206 |
| aromatic compound catabolic process | GO:0019439 | 196 | 5 | 0.7 | 7.11 | 0.0229 |
| heterocycle catabolic process | GO:0046700 | 190 | 5 | 0.68 | 7.34 | 0.0202 |
| nucleobase-containing compound catabolic process | GO:0034655 | 172 | 5 | 0.62 | 8.11 | 0.0137 |
| organophosphate metabolic process | GO:0019637 | 458 | 7 | 1.64 | 4.26 | 0.0385 |
| organophosphate catabolic process | GO:0046434 | 109 | 5 | 0.39 | 12.79 | 0.00215 |
| nucleoside phosphate catabolic process | GO:1901293 | 89 | 5 | 0.32 | 15.67 | 0.000958 |
| nucleoside diphosphate catabolic process | GO:0009134 | 69 | 5 | 0.25 | 20.21 | 0.000417 |
| glycolytic process | GO:0006096 | 63 | 5 | 0.23 | 22.14 | 0.000288 |
| muscle structure development | GO:0061061 | 66 | 3 | 0.24 | 12.68 | 0.0495 |
| muscle cell differentiation | GO:0042692 | 49 | 3 | 0.18 | 17.08 | 0.0227 |
| muscle cell development | GO:0055001 | 40 | 3 | 0.14 | 20.92 | 0.0139 |
| protein maturation | GO:0051604 | 234 | 6 | 0.84 | 7.15 | 0.0076 |
| protein folding | GO:0006457 | 132 | 6 | 0.47 | 12.68 | 0.000552 |
| *Unclassified* |  | 9777 | 20 | 35.06 | 0.57 | 0.0224 |

**Table 6.** PANTHER overrepresentation analysis of Molecular Function GO terms represented in the ANKB interactome. Fold enrichment is calculated from the observed number of genes/proteins in the ANKB interactome (# interactome) divided by the expected number of hits (expected), based on the total number of genes within the *Rattus norvegicus* genome classified under the same GO term (# genome). Enrichment was considered significant with a false discovery rate (FDR) < 0.05. Row shading indicates GO terms within a hierarchical set.

| PANTHER GO-Slim Molecular Function | GO ID | # genome | # interactome | expected | fold enrichment | FDR |
| --- | --- | --- | --- | --- | --- | --- |
| binding | GO:0005488 | 6194 | 45 | 22.21 | 2.03 | 9.33E-06 |
| protein-containing complex binding | GO:0044877 | 602 | 24 | 2.16 | 11.12 | 3.69E-16 |
| chromatin binding | GO:0003682 | 234 | 10 | 0.84 | 11.92 | 1.01E-06 |
| nucleosome binding | GO:0031491 | 67 | 10 | 0.24 | 41.63 | 6.33E-12 |
| organic cyclic compound binding | GO:0097159 | 2666 | 20 | 9.56 | 2.09 | 0.0449 |
| chromatin DNA binding | GO:0031490 | 72 | 10 | 0.26 | 38.74 | 1.08E-11 |
| protein binding | GO:0005515 | 2417 | 23 | 8.67 | 2.65 | 0.000583 |
| cytoskeletal protein binding | GO:0008092 | 465 | 17 | 1.67 | 10.2 | 6.3E-11 |
| actin binding | GO:0003779 | 225 | 15 | 0.81 | 18.59 | 8.24E-13 |
| actin filament binding | GO:0051015 | 167 | 13 | 0.6 | 21.71 | 6.08E-12 |
| unfolded protein binding | GO:0051082 | 54 | 4 | 0.19 | 20.66 | 0.00233 |
| lyase activity | GO:0016829 | 114 | 4 | 0.41 | 9.79 | 0.0287 |
| carbon-oxygen lyase activity | GO:0016835 | 45 | 3 | 0.16 | 18.59 | 0.0228 |
| hydro-lyase activity | GO:0016836 | 44 | 3 | 0.16 | 19.02 | 0.0229 |
| structural molecule activity | GO:0005198 | 550 | 10 | 1.97 | 5.07 | 0.00163 |
| *Unclassified* |  | 10317 | 20 | 36.99 | 0.54 | 0.00662 |
| molecular transducer activity | GO:0060089 | 1954 | 0 | 7.01 | < 0.01 | 0.0382 |
| signaling receptor activity | GO:0038023 | 1954 | 0 | 7.01 | < 0.01 | 0.0405 |

**Table 7.**
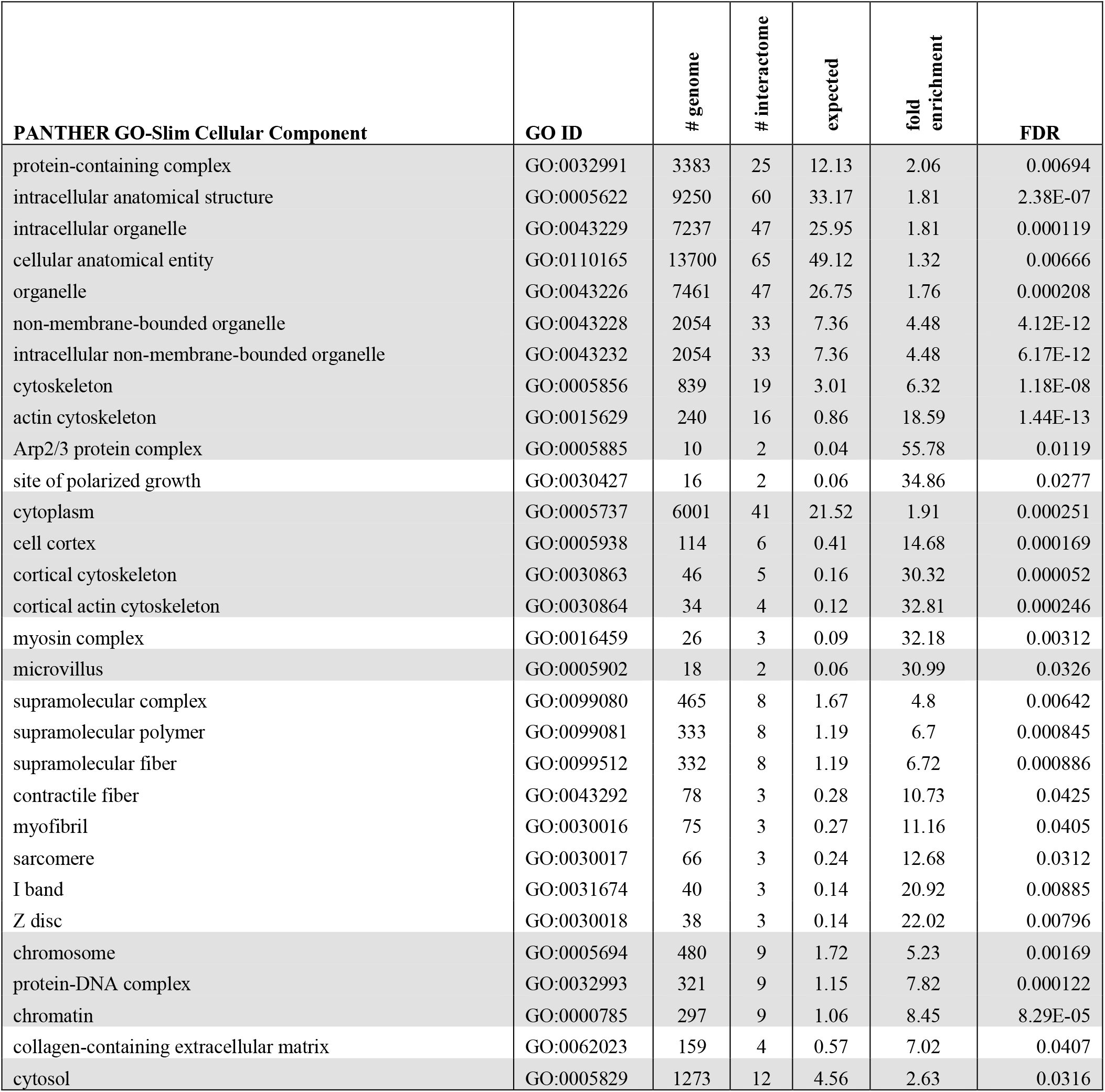
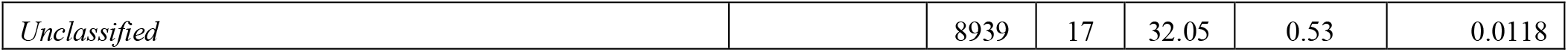
PANTHER overrepresentation analysis of Cellular Component GO terms represented in the ANKB interactome. Fold enrichment is calculated from the observed number of genes/proteins in the ANKB interactome (# interactome) divided by the expected number of hits (expected), based on the total number of genes within the *Rattus norvegicus* genome classified under the same GO term (# genome). Enrichment was considered significant with a false discovery rate (FDR) < 0.05. Row shading indicates GO terms within a hierarchical set.

**Table 8.** PANTHER overrepresentation analysis of PANTHER Protein Class terms represented in the ANKB interactome. Fold enrichment is calculated from the observed number of genes/proteins in the ANKB interactome (# interactome) divided by the expected number of hits (expected), based on the total number of genes within the *Rattus norvegicus* genome classified under the same PANTHER Protein Class term (# genome). Enrichment was considered significant with a false discovery rate (FDR) < 0.05. Row shading indicates GO terms within a hierarchical set.

| PANTHER Protein Class | PANTHER ID | # genome | # interactome | expected | fold enrichment | FDR |
| --- | --- | --- | --- | --- | --- | --- |
| chaperone | PC00072 | 225 | 7 | 0.81 | 8.68 | 0.000651 |
| Hsp90 family chaperone | PC00028 | 6 | 3 | 0.02 | > 100 | 4.34E-05 |
| Hsp70 family chaperone | PC00027 | 14 | 2 | 0.05 | 39.84 | 0.0221 |
| cytoskeletal protein | PC00085 | 648 | 18 | 2.32 | 7.75 | 7.78E-10 |
| actin or actin-binding cytoskeletal protein | PC00041 | 285 | 17 | 1.02 | 16.64 | 4.03E-14 |
| non-motor actin binding protein | PC00165 | 105 | 5 | 0.38 | 13.28 | 0.00125 |
| chromatin/chromatin-binding, or -regulatory protein | PC00077 | 327 | 15 | 1.17 | 12.79 | 6.41E-11 |
| lyase | PC00144 | 142 | 4 | 0.51 | 7.86 | 0.0307 |
| Unclassified |  | 7397 | 11 | 26.52 | 0.41 | 0.00348 |
| transmembrane signal receptor | PC00197 | 2071 | 0 | 7.43 | < 0.01 | 0.0161 |

**Table 9.** PANTHER overrepresentation analysis of PANTHER Pathways terms represented in the ANKB interactome. Fold enrichment is calculated from the observed number of genes/proteins in the ANKB interactome (# interactome) divided by the expected number of hits (expected), based on the total number of genes within the *Rattus norvegicus* genome classified under the same PANTHER Pathway term (# genome). Enrichment was considered significant with a false discovery rate (FDR) < 0.05. Row shading indicates GO terms within a hierarchical set.

| **PANTHER Pathways** | **PANTHER ID** | **# genome** | **# interactome** | **expected** | **fold enrichment** | **FDR** |
| --- | --- | --- | --- | --- | --- | --- |
| ATP synthesis | P02721 | 10 | 2 | 0.04 | 55.78 | 0.0181 |
| Glycolysis | P00024 | 56 | 4 | 0.2 | 19.92 | 0.00263 |
| Cytoskeletal regulation by Rho GTPase | P00016 | 82 | 5 | 0.29 | 17.01 | 0.000927 |
| Parkinson disease | P00049 | 99 | 4 | 0.35 | 11.27 | 0.018 |
| Integrin signalling pathway | P00034 | 189 | 5 | 0.68 | 7.38 | 0.016 |
| *Unclassified* |  | 20260 | 55 | 72.64 | 0.76 | 3.12E-05 |

Finally, we performed Kyoto Encyclopedia of Genes and Genomes (KEGG) pathway enrichment analysis [28,29] to further understand potential molecular interactions and reaction networks within our ANKB interactome. . Several pathways involved in cytoskeletal organization and metabolism were enriched, including regulation of the actin cytoskeleton (rno04810; +7.36 fold enrichment, FDR = 0.00155), cytoskeleton in muscle cells (rno04820; +11.14 fold enrichment, 4.71 x 10^-6^), tight junctions (rno04530; +8.29 fold enrichment, FDR = 0.00299), glycolysis/gluconeogenesis (rno00010; +24.42 fold enrichment, FDR = 5.04 x 10^-6^), and carbon metabolism (rno01200; +13.32 fold enrichment, FDR = 1.04 x 10^-4^). Together, our bioinformatics analyses support the participation of ANKB in cytoskeletal, chromatin-associated, metabolic, and muscle development and structure-associated networks.

## Discussion

Here, we presented an expanded cohort, now including 68 individuals with the ANKB p.S646F variant, building on our original report from 2017 [1]. Founder populations provide a unique opportunity to assess the clinical phenotype across the life course. Identifying additional individuals with the p.S646F variant consistently exhibited structural cardiac abnormalities including cardiomyopathy, and congenital heart anomaly. Although other manifestations were also apparent, such as sudden death, our study focused on the role of ANKB in sarcomere structure. Our cellular studies demonstrate that loss of ANKB expression disrupted normal sarcomere organization in primary cardiomyocyte cultures. Finally, our unbiased proteomics screen in H9c2 cells identified a novel interaction between ANKB and α-actinin-1, a focal adhesion-associated protein recently implicated in sarcomere organization during cardiomyocyte development [20], which was enhanced in the presence of the p.S646F variant. Bioinformatics analyses of our novel interactome strongly implicate ANKB in actin cytoskeleton regulation, providing a potential mechanistic basis for the defects we observed in sarcomere organization. In addition to cytoskeletal proteins, the ANKB interactome was enriched for chaperone, metabolic, and chromatin networks. Collectively, our findings support a previously understudied role for ANKB in regulating cardiomyocyte structure and suggest a model in which ANKB-dependent alterations in cytoskeletal organization contribute to the structural cardiac phenotypes associated with the p.S646F variant.

ANKB-associated cardiac diseases have historically been considered primarily electrical disorders, with pathogenic variants linked to a spectrum of heart rhythm disturbance [2–4]; however, accumulating evidence increasingly implicates ANKB in structural disease as well. Given the essential role for ANKB in targeting and anchoring diverse ion channels, transporters and receptors [6,7], the prominence of electrical phenotypes within the ANKB disease literature is perhaps unsurprising. Indeed, upon our initial identification of the ANKB p.S646F variant in two multigenerational Gitxsan First Nation families [1], we observed heart rhythm disturbance, seizures, and sudden cardiac death among those with the p.S646F variant. However, we also identified cardiac structural abnormalities within a subset, although the small initial cohort precluded assessment of their broader prevalence. Our expanded cohort further supports and extends upon our initial findings, demonstrating that p.S646F carriers indeed exhibit a range of cardiac structural abnormalities, including ventricular hypertrophy, dilated CMO, and congenital heart anomalies (Table 1). Consistent with our findings, structural cardiac phenotypes have been reported in association with other ANKB variants [5,30,31].

Our identification of a role for ANKB in sarcomere organization expands an understudied body of work implicating ANKB in the cardiac contractile apparatus. Previous studies demonstrated that ANKB localizes to the A-band, Z-line, and M-line of the sarcomere in cardiomyocytes [14,21], with localization dependent on the ANKB isoform [32]. For example, the striated muscle-specific ANKB-212 isoform interacts with obscurin, a large cytoskeletal scaffold and signalling hub, to establish its localization to the M-line [14,32,33]. Despite the localization of ANKB to sarcomere structures, a role for ANKB in sarcomere organization was not previously described. These prior studies examining ANKB function in cardiomyocytes employed isoform specific siRNA-mediated depletion and germline ANKB knockout models, and focused primarily on ANKB localization and associated membrane protein organization rather than sarcomere structure. In contrast, our study used cardiomyocyte-specific ANKB deletion using the Myh6-Cre driver, which is active in the embryonic and postnatal periods, and demonstrates that both partial and complete loss of ANKB disrupts sarcomere organization in primary neonatal cardiomyocytes. The effects of partial ANKB loss are of particularly interest in the context of human disease, in which p.S646F is heterozygous. Differences in the timing and cellular context of ANKB depletion across these models may influence the extent to which sarcomere organization is perturbed. This consideration is particularly relevant during cardiomyocyte development, when sarcomere structures undergo extensive remodelling and maturation [34]. Defining whether ANKB influences formation, maturation, or maintenance of these structures will require temporally controlled ANKB depletion approaches.

Using an unbiased proteomics approach in H9c2 cells, we identified and validated a novel interaction between ANKB and α-actinin-1, an actin-crosslinking protein, which increased approximately two-fold with the p.S646F variant despite unchanged total α-actinin-1 levels. This suggests that the variant may alter ANKB conformation, interaction site accessibility, or the composition of shared protein complexes. Given the emerging role of α-actinin-1 in coordinating focal adhesion maturation with cardiac sarcomere assembly [20], altered ANKB-α-actinin-1 binding may influence the organization or maturation of cytoskeletal structures that couple contractile machinery to sites of force transmission. Our proteomics screen also identified α-actinin-4, another non-muscle α-actinin reported to localize to the Z-disk and regulate sarcomere dynamics [35], as a candidate ANKB interactor, although this association has yet to be validated. Given the identification of multiple α-actinin family members as well as other cytoskeletal-associated proteins within the ANKB interactome, these findings suggest that the ANKB-α-actinin-1 interaction could represent one component of a broader cytoskeletal network through which ANKB contributes to cardiomyocyte structural organization. Future studies examining focal adhesion organization and ANKB-dependent regulation of α-actinin-1 localization or function will be important to determine how this interaction contributes to cardiomyocyte structure. Moreover, whether altered ANKB–α-actinin-1 binding directly contributes to the sarcomere disorganization observed following ANKB depletion remains to be determined.

We also probed whether the ANKB p.S646F variant alters the previously established ANKB-IP3R interaction [10,36,37]. Consistent with earlier reports from cardiomyocytes [10,37], we detected an ANKB-IP3R interaction in H9c2 cells expressing ANKB WT. Despite the localization of p.S646F within the ANKB membrane-binding domain and the previous mapping of the IP3R interaction to this domain, IP3R enrichment was not altered by p.S646F expression. This finding suggests that the variant does not grossly disrupt the ANKB region required for IP3R association and, together with the enhanced ANKB-α-actinin-1 interaction described above, suggests that p.S646F selectively alters ANKB protein interactions. Notably, total IP3R protein levels were significantly reduced in the presence of ANKB p.S646F. Given previous evidence that ANKB promotes IP3R stability and localization in cardiomyocytes [10,37], this reduction may be due to the lower overall ANKB p.S646F protein expression compared to WT ANKB, although additional effects on IP3R stability, trafficking, and degradation remain possible. Whether the reduced IP3R abundance contributes to the cardiac phenotypes associated with p.S646F remains unknown.

Beyond these validated interactions, bioinformatics analyses of our H9c2-derived ANKB interactome revealed a broader ANKB interaction landscape encompassing proteins across multiple cellular processes. In addition to cytoskeletal-associated proteins, the ANKB interactome was enriched for proteins involved in cellular metabolic processes and chromatin organization. PANTHER overrepresentation analysis identified enrichment of ANKB interactors in pathways related to glycolysis and carbon metabolism. Given the high metabolic demands of cardiomyocytes (reviewed in [38,39]), these associations raise the possibility that ANKB may contribute to the organization or localization of metabolic machinery within cardiomyocytes, potentially in proximity to the energetically demanding contractile apparatus. Consistent with this possibility, recent studies have demonstrated that ANKB contributes to mitochondrial organization and bioenergetic homeostasis in striated muscle. ANKB has been implicated in regulating mitochondrial fission and sarcoplasmic reticulum-mitochondria coupling [40], as well as in linking mitochondrial-associated complexes to the sarcolemma [41]. These observations are consistent with the enrichment of metabolic proteins found in our interactome. The identification of chromatin-associated proteins within our ANKB interactome was particularly unexpected. Although this finding has not yet been validated, it raises the possibility of an unrecognized association between ANKB and nuclear or chromatin-associated protein complexes. The contractile apparatus is physically coupled to the nucleus in cardiomyocytes, enabling mechanical signals generated at the sarcomere to be transmitted into the nucleus, influencing gene expression (reviewed in [42]). Given the structural link, ANKB could potentially contribute to this mechanotransduction network, although this possibility requires further investigation. Together, our findings expand upon the established view of ANKB as a membrane scaffold linking ion channels and transporters to the cytoskeleton. Our data suggest that ANKB additionally participates in a broader network of structural, metabolic, and nuclear-associated protein complexes within cardiomyocytes. Although ANKB WT and p.S646F samples were combined to define the broader ANKB interaction landscape, comparison of the two datasets revealed notable differences. Most strikingly, the p.S646F variant was associated with a greater number of interactors than ANKB WT. Among these were several molecular chaperones, including HSP90 and HSP70 family members, which facilitate protein folding, refolding and degradation [43]. This raises the possibility that the p.S646F variant exhibits altered protein folding or stability, and thus may have an increased dependence on chaperone-mediated stabilization compared to ANKB WT. This interpretation is further supported by our previous data [1,15], recapitulated here (Figure 2 C, D), demonstrating reduced p.S646F protein levels compared to ANKB WT following expression in H9c2 cells, consistent with decreased protein stability. These findings may be particularly relevant in the context of the age-dependent clinical phenotypes observed among p.S646F carriers in our cohort, including the relative scarcity of QTc prolongation among individuals below the age of 25. Within our cohort, individuals with cardiomyopathy ranged in age from 24 to 71 years, ranging from a 24-year-old male with DCM to a 71-year-old female with HCM. Chaperone activity declines with age [44,45], while the accumulation of misfolded and damaged proteins increases [43]. Thus, an increased dependence of ANKB p.S646F on chaperone-mediated folding or stabilization could render the variant increasingly vulnerable to age-related declines in proteostasis. Alternatively, increased chaperone engagement by the variant may contribute to an overall competition for chaperone capacity within the cell. Together, it is reasonable to suggest that age-related changes in proteostasis could promote the emergence or progression of cardiac phenotypes in ANKB p.S646F carriers.

Collectively, our findings extend the established role of ANKB beyond the organization of membrane-associated ion channels and transporters. In addition to demonstrating a requirement for ANKB in sarcomere organization, our unbiased proteomics analysis identified candidate interactions across cytoskeletal, metabolic and chromatin-associated proteins, suggesting that ANKB contributes to the spatial organization of multiple structural and signalling domains within cardiomyocytes. Although many of these putative interactions were identified in H9c2 cells and require validation in primary cardiomyocytes and intact tissue, they provide a framework for future studies. Complementary biochemical and imaging approaches, such as immunostaining and proximity ligation assays, will help establish their subcellular localization and functional significance.

The relationship between our structural findings and the established roles of ANKB in ion channel organization and cardiomyocyte excitability also warrants further investigation. Rather than representing independent functions, cardiomyocyte structure and function are likely to be closely interconnected. Altered cytoskeletal organization could influence ion channel localization, calcium handling, and excitation-contraction coupling, while perturbations in these processes may themselves promote structural remodelling. Our findings of a structural role for ANKB are therefore unlikely to be independent of its well-established role in regulating cardiac excitability. Moreover, whether the cellular structural abnormalities identified here contribute directly to the tissue-level remodelling observed in ANKB-associated cardiac disease, including the structural cardiac abnormalities observed in p.S646F carriers, remains to be determined. Defining the temporal and mechanistic relationships between these processes will be an important direction for future investigation.

## Methods

### Clinical assessment

Participants were recruited as part of a larger study in Northern British Columbia First Nations populations [46]. Individuals were invited to enroll if they had clinical features of LQTS, a personal or family history of arrhythmia or sudden death, or were related to an individual carrying the ANKB c.1937C>T (p.S646F) variant. Community approval and individual informed consent were obtained under protocols following University of British Columbia (UBC H05-70330-A073) and Northern Health Authority (RRC-2007-0038) ethics approvals. Referrals occurred through affected family members, cascade screening, and healthcare providers. Clinical information was collected through questionnaires and medical records. Participants were also tested for the known Northern British Columbia First Nations variants including the LQTS-associated KCNQ1 variants p.V205M [46] and p.R591H [17], as well as the p.L353L modifier and the KCNH2 modifier p.K897T [47]. Full data collection was initiated in 2004, reported on in [46,47], and specifically for the S646F variant, in [1]. Data collection continued until March 2026 and was compiled by AM, for the purpose of this study.

In short, cardiac assessment was conducted through the adult and pediatric BC Inherited Arrhythmia Program (BCIAP) clinics when available. Clinical assessments included electrocardiograms (ECGs), exercise stress testing, Holter monitoring, and echocardiography. Clinical features reviewed included QTc prolongation, arrhythmias, syncope, seizures, structural cardiac abnormalities, and sudden death. All ECGs on file were reviewed by cardiac electrophysiologists blinded to the variant and clinical status. The QT interval was measured manually using the tangent method [48]. The reported QT interval was the longest identified within all 12 leads and corrected for heart rate using the Bazett formula. All data was collected and compiled within the larger LQTS project (AM).

For sudden deaths, family members were spoken to regarding circumstances by (AM or LA), coroners’ reports and autopsies were accessed with consent and study information, such as genotype was documented.

### H9c2 cell culture

The H9c2 rat ventricular-derived cardiomyoblast cell line was obtained from the American Type Culture Collection (ATCC, CRL-1446). H9c2 cells were cultured in Dulbecco’s Modified Eagle Medium (DMEM, ATCC 30-2002) supplemented with 10% fetal bovine serum (FBS, Thermo Fisher, 12483020, reserve 10498874, lot 2240764RP), 100 U/mL penicillin, and 100 μg/mL streptomycin at 37°C in 5% CO2. Cells were kept at or below 15 passages and passaged at 70-80% confluence. Each vial was tested for mycoplasma using the universal mycoplasma detection kit (ATCC, 30-1012K). One day after seeding, cells were transfected with either GFP control, ANKB WT-GFP, or ANKB S646F-GFP using jetPEI (Sartorius), according to the manufacturer’s instructions. Briefly, 4 µL of jetPEI diluted in 150 mM NaCl was added to 2 µg of plasmid DNA also diluted in 150 mM NaCl, the mixture was incubated for 15 minutes and then added dropwise to the cells in DMEM containing no antibiotics.

### Immunoprecipitations

GFP control, ANKB WT-GFP, or ANKB S646F-GFP expressing H9c2 cells were collected 48 hours following transfection. First, cells were washed twice with TBS. Then approximately 4.4×10^7^ cells per condition were homogenized on ice in RIPA buffer supplemented with protease inhibitor cocktail for 30 minutes, followed by centrifugation at 4°C for 20 minutes at 12,000 rpm to remove debris. Supernatants were transferred to fresh tubes, and protein concentration was determined using a modified Lowry assay. For each immunoprecipitation, 50 µL Dynabeads Protein G were washed and incubated with 5 µg antibody diluted in 200 µL TBS-T (0.02% Tween-20, pH 7.4) for 30 minutes at 4°C with rotation. Antibody-bound beads were washed once with TBS-T, washed twice with conjugation buffer (150 mM NaCl, 50 mM Tris-HCl, pH 7.4), and cross-linked by incubation in 250 µL 5 mM BS3 for 30 minutes at room temperature with rotation. Cross-linking was quenched with 12.5 µL quenching buffer (1 M Tris-HCl, pH 7.5) for 15 minutes at room temperature, and beads were washed three times with RIPA/TBS-T. Prepared antibody-bead complexes were incubated with 2 mg lysate for 2 hours at 4°C with rotation. Beads were washed twice with RIPA buffer and once with TBS-T, then eluted in two bead volumes of 0.5 M ammonium hydroxide/0.5 mM EDTA for 30 minutes at room temperature with rotation. Samples were then split into 4 microcentrifuge tubes and dried by SpeedVac. One microcentrifuge tube was reconstituted in RIPA buffer and mixed with SDS-PAGE loading dye under reducing conditions (dithiothreitol and β-mercaptoethanol) for Western blot analysis. The remaining sample was analyzed for protein interaction partners at the UVic-Genome BC Proteomics Centre, Victoria, Canada, using high performance liquid chromatography coupled to tandem mass spectrometry (LC-MS).

### Trypsin Digestion

The following procedures were performed at the UVic-Genome BC Proteomics Centre. Immunoprecipitation samples were rehydrated in 9 M urea/300 mM Tris pH 8.0, and protein concentration was determined by bicinchoninic acid (BCA) assay. 4 µg of protein was reduced with dithiothreitol (DTT) to a final concentration of 20 mM and incubated at 37°C for 30 minutes. Samples were alkylated with iodoacetamide to a final concentration of 40 mM for 30 minutes at room temperature in the dark. Samples were diluted with 100 mM Tris pH 8.0 to reduce the urea concentration to 0.78 M and digested at 37°C for 18 hours with modified sequencing grade porcine trypsin (Promega) at a 50:1 substrate-to-enzyme ratio. The samples were acidified with formic acid to a final concentration of 0.99% and desalted and concentrated using Pierce C18 Tips (Thermo Scientific) following manufacturers protocols. The peptides were then eluted in 70% acetonitrile/0.1% formic acid and dried under nitrogen gas flow. Samples were rehydrated in 25 mM ammonium bicarbonate and further cleaned using Pierce Detergent Removal Spin Columns (Thermo Scientific) according to the manufacturer’s instructions. Detergent-free samples were frozen, lyophilized overnight, and rehydrated in water/0.1% formic acid to make 0.025 µg/µL prior to LC-MS analysis.

### Liquid chromatography and tandem mass spectrometry (LC-MS)

Based on quantitation values, 0.5 µg of each sample was loaded onto a Evotip Pure™ Tip following the manufacturer’s protocol. Peptides were separated using an Evosep One system (Evosep Biosystems, Odense, Denmark) with an Evosep EV1106 C18 column (15 cm × 150 µm, 1.9 µm particle size, 30 µm ID stainless steel emitter tip) using the Xcalibur 15 SPD method (88-minute gradient). The chromatography system was coupled on-line with an Orbitrap Fusion Tribrid mass spectrometer (Thermo Fisher Scientific, San Jose, CA) equipped with a Nanospray Flex NG source. Solvents were A: water with 0.1% formic acid; and B: 100% acetonitrile with 0.1% formic acid.

The Orbitrap Fusion instrument parameters (Fusion Tune 4.2.4321 software) were as follows for orbitrap (OT-MS) iontrap (OT-MS/MS) with HCD fragmentation: spray voltage 1950 V and capillary temperature 300°C. Survey MS1 scans were acquired over a mass range of 350–1500 m/z in profile mode at resolution 120,000 FWHM at 200 m/z one microscan with Automatic inject time with the Siloxane mass 445.12002 used as lock mass for internal calibration. Data-dependent acquisition scheduled Orbitrap survey scans every 3 seconds, with the software determining “Top number most intense” ions for MS/MS acquisitions during this period. The automatic gain control (AGC) target value for FTMS was set to Standard 100% (400,000 counts) and automatic maximum fill time to ensure optimal sensitivity and cycle time. The most intense ions charge state 2-5 exceeding 25,000 counts were selected for HCD MSMS fragmentation in the ion routing multipole. Monoisotopic precursor selection most abundant peptide peak was enabled, and Dynamic exclusion settings were repeat count: repeat count 2, repeat duration 5 seconds, exclusion duration 7.5 seconds with a 10 PPM mass window. The data dependent (ddMS2) IT HCD scan used a quadrupole isolation window of 1.6 Da; Iontrap rapid scan rate, automode normal m/z range, centroid detection, 1 microscan, Auto maximum injection time, AGC target (200%) or 20,000 counts and stepped HCD collision energy of 30 and 35%.

### Data analysis parameters

Tandem mass spectra were extracted and deisotoped by Proteome Discoverer version 3.0. Charge state deconvolution was not performed. All MS samples were analyzed using Sequest (Thermo Fisher Scientific, San Jose, CA, USA; version IseNode in Proteome Discoverer 3.0.0.757). Sequest was set up to search contaminants.fasta; and *Rattus norvegicus*, assuming the digestion enzyme trypsin. Sequest was searched with a fragment ion mass tolerance of 0.60 Da and a parent ion tolerance of 10.0 PPM. Carbamidomethyl of cysteine was specified in Sequest as a fixed modification. Oxidation of methionine, acetyl of the n-terminus were specified in Sequest as variable modifications. The datasets from three individual biological replicates were combined, and proteins identified in GFP control samples were removed from the final ANKB interaction partners list.

### Protein interaction analysis

The curated list of ANKB protein interaction partners identified by LC-MS was analyzed using STRING to identify a protein-protein interaction network [25]. The resulting interactions were then graphed in R (version 4.4.0). Specific Gene Ontology (GO) terms were extracted from ShinyGO to highlight specific protein-protein interaction networks. The ANKB interactome was further analyzed for functional enrichment using GO and Kyoto Encyclopedia of Genes and Genomes (KEGG) pathway analyses. GO enrichment analysis was performed using the Gene Ontology Resource (release 2026-03-25) [49,50] and PANTHER enrichment analysis (version 19.0) [27] to identify overrepresented biological processes, molecular functions, and cellular components among the ANKB interactome. An overrepresentation test was conducted using a Fisher’s Exact Test with a Benjamini-Hochberg False Discovery Rate (FDR) correction for multiple testing. A corrected p-value (or FDR) of <0.05 was considered statistically significant. The analysis was conducted against a reference list of all *Rattus norvegicus* genes available in the PANTHER database. GO-Slim classification was performed for Biological Process, Molecular Function, and Protein Class categories to determine the proportion of interacting proteins within each group. Enrichment results were graphed in GraphPad Prism (version 11.0.0). KEGG pathway enrichment analysis was performed using ShinyGO (version 0.82) and then visualized in R (version 4.4.0).

### Experimental animals

All animal procedures were approved by the University of Victoria Animal Care Committee and performed in accordance with the guidelines set by the Canadian Council on Animal Care. AnkB^flox/flox^ mice were a kind gift from Dr. Peter Mohler (Ohio State University) and were back-crossed in-house for 5 generations onto a C57BL/6J background (000664, The Jackson Laboratory, [51]). AnkB^flox/flox^ mice were crossed with Myh6-Cre (#011038, The Jackson Laboratory, [52]) to generate control, heterozygous knockout (HET), and knockout (KO) littermates. The αMyHC promoter is transcriptionally active in the embryonic period [53], with a marked increase in activity shortly following birth, in part due to the rise in thyroid hormone expression [52]. Mice were housed under a 12/12-hour light/dark cycle starting at 8 A.M., with food and water ad libitum; temperature was maintained between 20°C and 25°C and humidity at 40–65%.

### Single heart primary cardiomyocyte cultures

Primary cardiomyocytes were isolated from postnatal day 1–3 (P1–P3) ANKB control, HET, and KO mouse hearts. Heart tissue was minced into fragments and incubated overnight at 4°C in isolation medium consisting of HBSS supplemented with 20 mM 2,3-butanedione monoxime and 0.0125% trypsin with gentle agitation. Following overnight digestion, tissue was incubated with L-15 medium supplemented with 20 mM butanedione monoxime and digestion medium containing 0.75 mg/mL collagenase/dispase and 20 mM butanedione monoxime for 30 minutes with gentle agitation at 37°C. Cells were mechanically dissociated with a serological pipette containing a narrowed tip. Tissue that was not dissociated in this step underwent further digestion with another incubation in digestion medium containing 0.75 mg/mL collagenase/dispase and 20 mM butanedione monoxime in L-15 medium by incubating at 37°C with gentle agitation for 5-10 minutes followed by more mechanical dissociation. Cells were centrifuged at 200 x g for 5 minutes and then resuspended in plating media. Plating media was supplemented DMEM with 5% heat-inactivated horse serum and 10% heat-inactivated FBS. Cells were plated at 1.4 x 10^5^ cells/cm^2^. Note, cells prepared for western blotting first were replated for 1.5-2 hours and then plated at 1.4 x 10^5^ cells/cm^2^. After 18-24 hours, cells were adhered and underwent a media change into maintenance media, DMEM supplemented with 5% heat-inactivated horse serum and penicillin/streptomycin and maintained for 2-3 days before being collected for immunostaining or Western blotting.

### Immunocytochemistry

Primary cardiomyocytes were seeded onto collagen coverslips either purchased from Neuvitro (NEU-GG-12-1.5-COLLAGEN) or coated in lab using collagen from calf skin (Fisher scientific, 1705061). Coverslips were fixed with 4% paraformaldehyde for 10 minutes, followed by three washes with PBS containing Ca^2+^ or Mg^2+^. Cells were permeabilized with 0.1% Tween-20 for 5 minutes, washed three times in PBS as before, blocked with 1% BSA and 22.52 mg/mL glycine in PBS-T, and incubated in α-actinin (1:400, Sigma, A7811) diluted in 1% BSA in PBS overnight at 4°C. Following primary antibody incubation, coverslips were washed three times in PBS for 5 minutes and then incubated with Hoechst (1:500) and secondary antibody (1:400, 488 Donkey anti-mouse, Jackson Immuno Research, 715-545-151) diluted in 1% BSA in PBS at room temperature for 1 hour. Following secondary incubation, coverslips were washed three additional times and mounted with VECTASHIELD (Vector Labs) and sealed using Sally Hansen Insta-dri nail polish.

### Microscopy

Cells were imaged using a Leica TCS SP8 confocal microscope with PMT detectors. Images were acquired using a 40× oil-immersion objective (1.30 NA, pinhole = 65.08 μm, 2048 × 2048 (pixel size = 141.98 nm × 141.98 nm)). GFP was excited using the 488 nm Argon laser at 20% intensity. For all images presented, brightness and contrast were uniformly adjusted for presentation purposes.

### Image analysis

Sarcomere organization was quantified using the previously published SarcOmere Texture Analysis (SOTA) method [19]. Images were analyzed in MATLAB using Haralick texture features derived from gray-level co-occurrence matrices to assess sarcomeric periodicity and organization. Prior to analysis, α-actinin images were filtered using a 3 × 3 median filter to reduce noise and then sharpened using the imsharpen function with a radius of 2 and amount of 1.5. The sarcomere organization score was then averaged across ROI for each replicate and then graphed and analyzed in GraphPad Prism.

### Western Blotting

Adherent cells (both transfected H9c2 and primary cardiomyocytes) were washed 2 times in dPBS and lysed in RIPA lysis buffer (9.1 mM PBS, 150 mM NaCl, 1.7 mM monobasic sodium phosphate, 9.1 mmol/L dibasic sodium phosphate, 1% Igepal, 0.5% w/v sodium deoxycholate, 0.1% sodium dodecyl sulphate (SDS), 10 μL/mL protease inhibitor cocktail (Sigma, P8340-5ML)) by scraping cells off the plate on ice. Protein lysates were pushed through a 27 ½ gauge needle twice and incubated on ice for 30 minutes; samples were then centrifuged at 13500 xg for 20 minutes at 4 °C. Protein content was determined using the DC protein assay (Bio-rad, 5000112), then 10-15 μg of protein was incubated in Laemmli sample buffer (1×: 0.08 mM bromophenol blue, 0.05 M Tris-HCl pH 6.8, 0.058 M SDS, 3.33% v/v glycerol) at room temperature under reducing conditions (5% dithiothreitol (DTT) and 5% β-mercaptoethanol), separated by SDS-PAGE on TGX-stain free gels (Bio-rad, 4561094) and transferred to a 0.2 μm pore PVDF membrane (Bio-rad, 1620177). The membrane was blocked with 5% fat free skim milk in TBS-T and then incubated with indicated antibodies in the blocking solution, and protein bands were visualized using the Clarity Western enhanced chemiluminescence substrate (Bio-Rad, 1705061) and a Syngene G:Box imager. All proteins were normalized to total protein using Ponceau S Solution (0.1% w/v in 5% acetic acid) (Sigma, P7170-1L) and quantified in ImageJ. The antibodies and concentrations used are listed in Table S1. Graphs were created in GraphPad Prism.

### Statistical analysis

All statistical analyses were performed using GraphPad Prism 11.0. Relevant details are described in Results and figure legends. Data are reported as mean ± standard error of the mean (SEM). Significance thresholds were defined as follows: ****,p < 0.0001; ***,p < 0.001; **,p < 0.01; *,p < 0.05; and non-significant (ns) for p > 0.05. Unpaired t-tests and two-way ANOVAs with Šídák’s multiple comparisons tests were performed as indicated. Assumptions of normality and homogeneity of variance were visually assessed using Q–Q plots and homoscedasticity plots of model residuals, respectively. These plots did not show substantial deviations from normal distribution.

## Abbreviations

ANKB: Ankyrin-B
NKA: sodium potassium ATPase
NCX: sodium calcium exchanger
IP3R: inositol 1,4,5-trisphosphate receptor
PP2A: protein phosphatase 2A
RYR2: ryanodine receptor 2
LQTS: long QT syndrome
VGCCs: voltage-gated calcium channels
GO: Gene Ontology
KEGG: Kyoto Encyclopedia of Genes and Genomes
DMEM: Dulbecco’s Modified Eagle Medium
FBS: fetal bovine serum
TBS: Tris-buffered saline
RIPA: radioimmunoprecipitation assay buffer
TBS-T: Tris-buffered saline + 0.2% Tween-20
BS3: bis(sulfosuccinimidyl) suberate
EDTA: ethylenediaminetetraacetic acid
LC-MS: liquid chromatography tandem mass spectrometry
BCA: bicinchoninic acid
DTT: dithiothreitol
HCD: higher-energy collisional dissociation
AGC: automatic gain control
FWHM: full width at half maximum
PPM: parts per million
HET: heterozygous knockout
KO: knockout
αMyHC: alpha myosin heavy chain
ICC: immunocytochemistry
PBS: phosphate-buffered saline
BSA: bovine serum albumin
WGA: wheat germ agglutinin
PMT: photomultiplier tube
NA: numerical aperture
ROI: region of interest
SOTA: SarcOmere Texture Analysis
SDS: sodium dodecyl sulfate
co-IP: co-immunoprecipitation
STRING: Search Tool for the Retrieval of Interacting Genes/Proteins
ACTN1/4: alpha-actinin-1/ alpha-actinin-4
Cav1.2: voltage-dependent L-type calcium channel subunit alpha-1C
PLCE1: phospholipase C epsilon 1
Cx43: connexin-43
HSP70: heat shock protein 70
cTnT: cardiac troponin T
Wnt: Wingless-related integration site signaling pathway

## Acknowledgements

This work was supported by a Canadian Institutes for Health Research Project Grant (CIHR; PJT-169064) awarded to LTA and LAS as well as from CIHR grants (PJT-185887, PJT-189953) awarded to LAS. NY was supported by a Vanier Canada Graduate Scholarship (Canadian Institutes of Health Research Vanier) and University of Victoria Donor Awards. We wish to acknowledge the UVic-Genome BC Proteomics Centre, Victoria, Canada for mass spectrometry data acquisition and processing. We acknowledge that this research was conducted on the traditional territory of the Lək̓ʷəŋən (Songhees and Xwsepsəm/Esquimalt) Peoples, and the Lək̓ʷəŋən and W̱SÁNEĆ Peoples whose historical relationships with the land continue to this day. We also acknowledge the Gitxsan peoples and their leadership as this work was initiated for and is foundational for our ongoing research partnership on the mechanisms of inherited cardiac disease. The progress of this work has been presented at in-person community gatherings, and virtually by NY.

## Author contributions

NY, LEWS, LA, LAS conceptualized the manuscript. NY conducted all experiments with MP and HM assisting in Western blotting and imaging, KWLF in primary culturing and imaging. NY and CL performed analysis and statistics. ACHM compiled the clinical information. NY and KWLF created figures. NY, LEWS, LA, and LAS wrote and edited the manuscript.

## Data availability statement

The datasets generated during and/or analysed during the current study are available from the corresponding author on reasonable request.

## Additional information

